# Saccadic suppression enhances saliency of ecologically relevant stimuli in the optic tectum

**DOI:** 10.1101/2025.02.21.639462

**Authors:** Giulia Soto, Carina Thomas, Louis Funk, Tim C. Hladnik, Florian A. Dehmelt, Ziad M. Hafed, Aristides B. Arrenberg

**Affiliations:** Werner Reichardt Centre for Integrative Neuroscience, University of Tübingen, Tübingen, Germany; Hertie Institute for Clinical Brain Research, University of Tübingen, Tübingen, Germany

## Abstract

Saccadic suppression, a reduction in visual sensitivity around the time of rapid eye movements, is well-documented in primate psychophysics but its mechanisms and functions remain debated. Here, we use zebrafish to trace the origins of saccadic suppression and we demonstrate how saccadic suppression selectively enhances the visual salience of ecologically relevant stimuli. Saccadic suppression thus contributes to more than just compensation for rapid saccade-induced image shifts. We first established a behavioral correlate of saccadic suppression in larval zebrafish escape behavior. Then, using electrophysiology, we show that retinal ganglion cells jumpstart saccadic suppression in a spatial frequency dependent manner. Calcium imaging, combined with 360° visual stimulation and behavioral tracking, revealed that motor signals enhance peri-saccadic suppression strength in the optic tectum, where suppression lasts for more than 3000 ms. Notably, saccadic suppression is much weaker and more short-lived for stimuli related to hunting or escape behavior than for behaviorally less relevant global flashes. This unequal attenuation effectively increases the salience of the ecologically relevant stimuli in the optic tectum after saccades. Our results demonstrate that saccadic suppression integrates visual and motor signals to optimize sensory processing within neural constraints, and they provide insights into evolutionarily conserved visual strategies.

## Introduction

Saccadic eye movements are an efficient way of rapidly sampling the visual environment, employed by many species across the animal kingdom. However, by moving the eyes at several hundred degrees per second from one direction of gaze to another, visual flow is greatly disrupted, a disruption that we are blissfully unaware of in our daily perception. From a perceptual standpoint, this can be explained through a phenomenon known as saccadic suppression. Saccadic suppression describes a brief peri-saccadic reduction in perceptual sensitivity, and it has been investigated in primates for decades [1–5]. Physiological studies furthermore showed that individual neuronal responses in visual areas are concomitantly reduced peri-saccadically [5], and that responses to stimuli of lower spatial frequency are suppressed more strongly [3,6]. While the benefit of suppressing visual processing during saccades intuitively makes sense, its computation in the brain as well as the reasons underlying variable suppression strengths under different image conditions are still unclear.

Zebrafish (*Danio rerio*) offer the opportunity to investigate neural correlates of saccadic suppression across the visual system in a vertebrate “blueprint” brain without a visual cortex. Larval zebrafish start exhibiting spontaneous saccadic eye movement prior to 4 days post-fertilization (dpf); by 5 dpf, saccades are well coordinated and comparable to those of adults [7]. Moreover, brainstem control of saccadic eye movements is highly conserved across evolutionarily distant species like zebrafish and primates [8]. Finally, the optic tectum (OT) of zebrafish is homologous to the primate superior colliculus (SC), and both of them are highly sensitive to visual stimulation [9]. Thus, zebrafish are well suited to answer the long-standing debate in the field on the origins and purposes of saccadic suppression.

Considering the frequent occurrence of saccadic eye movements, it might be naïve to think that saccadic suppression simply serves the purpose of suppressing a visual disturbance. Suppression of visual information for fractions of a second with every executed saccade seems simply like too large of a loss of sensory information, particularly in a system strapped for neural resources, such as the small larval zebrafish brain. A potential purpose of saccadic suppression could, instead, be as a precise tool for efficient task-specific encoding. The observation that visual stimulus properties affect suppression dynamics would be in line with this possibility. For example, saccades across low spatial frequency surroundings will lead to stronger and longer lasting suppression, compared to high spatial frequency surroundings [6,10–12]. Similar effects can be observed with luminance contrast dependency [6,13] and chromatic contrast [14]. Thus, saccadic suppression can be highly specific and depends on image content. As we show here, the stimulus-dependence of saccadic suppression could even be beneficial to optimize sensory encoding in behavioral contexts, such as prey capture, where saccadic eye movements play a vital role in the behavioral sequence [15].

The preponderance of evidence for visual dependencies of saccadic suppression both illuminates and complicates the search for the circuit origins of saccadic suppression. For example, is saccadic suppression purely visual? Masking, or the reduced visibility of one brief visual stimulus due to the presence of a second brief visual stimulus [16], plays a vital role in generating saccadic suppression. In the 1970’s, there was already behavioral evidence that visual masking leads to the reduced perception of a “blur”, a visual disturbance caused by the saccadic eye movement itself [17,18]. More recently Idrees and colleagues showed in isolated mammalian retinae, that visual masking leads to reduced responses to a probe flash in retinal ganglion cells, following a saccade-like texture displacement [6,19]. However, saccadic motor signals still likely play a role. For example, it is known from psychophysics studies that the peri-saccadic window of suppression was significantly shortened when participants performed real saccades instead of watching a simulated corresponding visual stimulus (a texture displacement) [6]. Corollary discharge signals originating from motor commands can act as modulatory signals for visual neurons during rapid eye movements [4,20]. Recent work by Ali and colleagues has shown that for locomotion, a corollary discharge signal can in fact be found in the zebrafish OT, the sensory structure responsible for processing most visual inputs in the zebrafish brain [21]. In primates, behavioral and neuronal evidence suggest the presence of a saccade-related corollary discharge signal in parts of the visual system [22–25].

It is evident that both visual and motor-related signals shape saccadic suppression properties. However, how the two act together on the individual neuronal level to modulate visual responses is poorly understood. Here, we show in larval zebrafish how visual responses of tectal neurons are modulated by sensory and motor-related signals during peri-saccadic suppression. We do so, by imaging large populations of OT neurons through calcium imaging, while simultaneously using 360° whole-field visual stimulation and behavioral tracking in tethered animals. We find that both visual and motor related processes shape saccadic suppression differentially in the OT. The OT receives purely visually suppressed signals from the retina and further modulates them. Motor-related signals in the OT enhance saccadic suppression. Furthermore, stimuli related to hunting and escape behavior appear more salient, because they are less suppressed than behaviorally irrelevant global flashes, indicating optimized signal processing.

## Results

### Visually driven saccadic suppression exists in the zebrafish

To enable our investigation of peri-saccadic stimulus encoding, we first need to demonstrate that saccadic suppression does exist in zebrafish. Classically, saccadic suppression has been studied as a behavioral phenomenon [26]. Therefore, we initially tested whether zebrafish exhibit saccadic suppression in a behavioral essay. Free-swimming larval zebrafish were placed into a custom-built circular arena and exposed to visual stimulation from four horizontally placed LCD screens (Fig. 1A). The screens displayed a binary gaussian blur texture that was rapidly displaced horizontally to simulate saccadic eye movements. Texture displacements were paired with a global dark flash, which in larval zebrafish can evoke O-bend escape responses [27–30]. When the global dark flash was shown within 500ms prior up to 2000ms after texture displacement onset, the fish showed a lower escape probabilities (Fig. 1D), indicating that visual processing of the global dark flash is suppressed around the time of a saccade-like texture displacement. Response latencies (mean ± SD) were highly variable and not affected by texture displacements (response latency from dark flash onset = 962.2 ±408.4 ms, Fig. 1C). After having found behavioral evidence of saccadic suppression in zebrafish, we were prompted to investigate the neural origins of saccadic suppression in zebrafish. We started with recording in the retina, because this is the location where saccadic suppression first occurs in the mammalian visual system [6]. Thus, we isolated adult zebrafish retinae and performed multi-electrode array (MEA) recordings of retinal ganglion cells (RGCs) during visual stimulation (Fig. 2A). In order to simulate the visual transient of a saccade, retinae were shown a coarse or fine texture which was rapidly displaced in a saccade-like manner (Fig. 2B, left). Texture displacements generated a visual response in RGCs as can be seen in Fig. 2C. To test if saccade-like texture displacements could suppress responses of a secondary visual stimulus, at various time points in close temporal proximity to the texture displacement, a 17 ms long dark or bright whole-field flash was shown (Fig. 2B, right, Fig. 2D). Flash responses, when shown in temporal isolation (“Baseline”) were robust and in most cases much larger than texture displacement responses. In order to isolate flash-associated responses around the time of the texture displacement, texture displacement-associated responses were subtracted from flash responses as shown in Fig. 2D (second and third panel) as in previous studies [6]. After removal of this “saccade”-associated response it can be seen that flash responses are significantly suppressed around the time of the texture displacement (Fig. 2D, third panel). When averaging all RGCs recorded (n=87), it became apparent that suppression is strongest at 17 and 50 ms after texture displacement onset and that responses return to baseline levels after ∼300 ms (Fig. 2E). This is a similar time course to that described in mouse and pig retinae [6]. Overall suppression strength is, however, weaker in zebrafish, with a maximal suppression of 15.8 ± 3.0% (mean ± SEM), whereas mammalian retinal saccadic suppression can be as strong as ∼30% (see Ref. 6, Fig. 3). Furthermore, when comparing the suppression induced by a coarse texture displacement versus a fine texture displacement, coarse will generally lead to stronger and longer lasting suppression, than fine, reproducing spatial frequency dependencies of saccadic suppression in primates [3,6,10].

**Fig 1.**
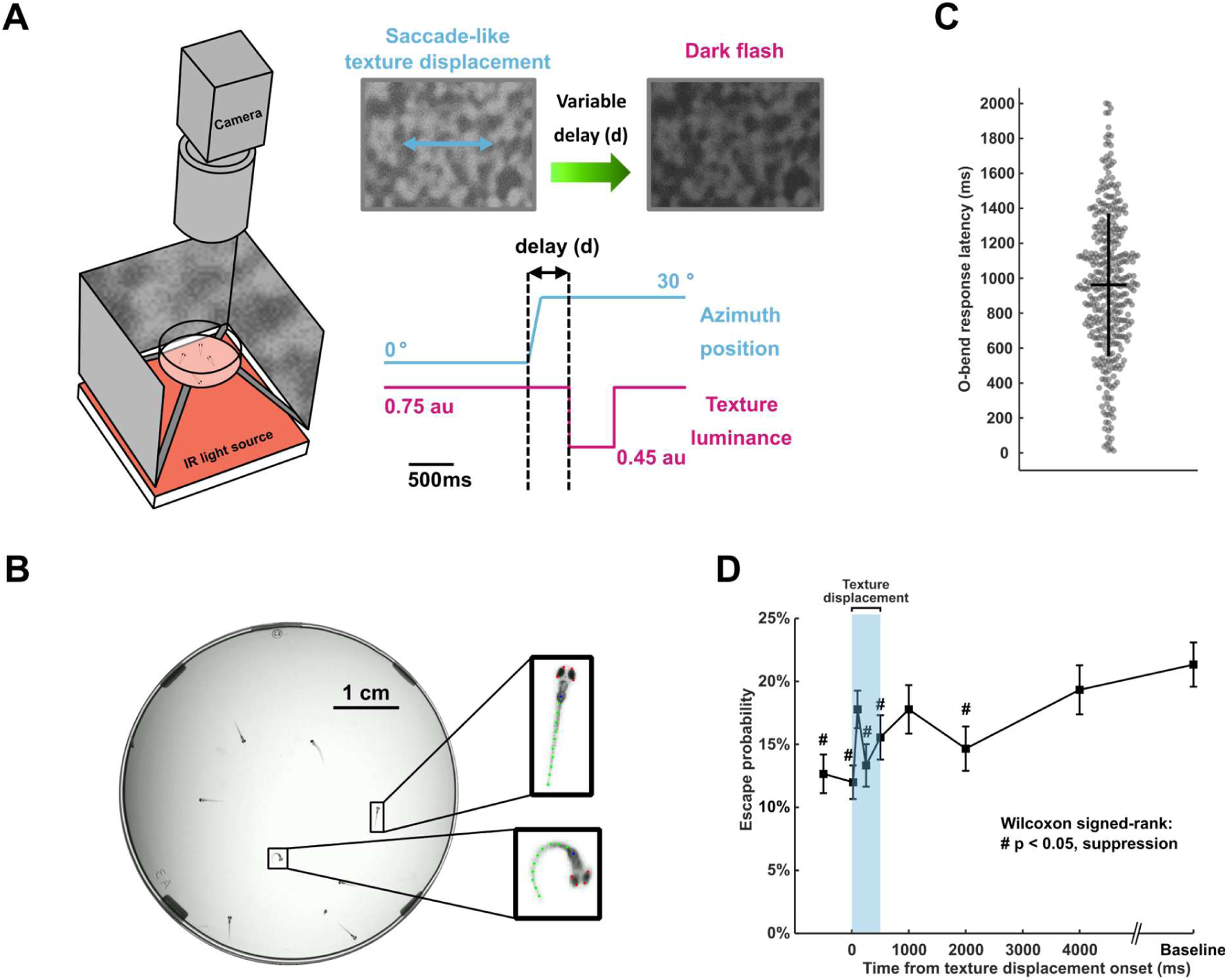
Larval escape behavior is affected by saccadic suppression. (**A**) Escape behavior (re-orienting O-bends) of free-swimming larval zebrafish was evoked by a dark flash and measured around the time of a saccade-like texture displacement (coarse texture). (**B**) In each recording 10 fish were tracked. Each fish was annotated with 15 landmarks across the eyes, swim bladder and tail. O-bends events were detected for tail bends >130°. (**C**) Escape latency from dark flash onset (962.2 ±408.4 ms, mean ± SD) was highly variable and not affected by texture displacements. (**D**) Escape probabilities (mean ± SEM) were reduced around the time of a texture displacement, compared to a baseline condition, where dark flashes were shown in the absence of a dark flash. (# p < 0.05, two-tailed Wilcoxon signed-rank test, Bonferroni corrected for n= 8 conditions, *n = 45 fish*).

**Fig. 2.**
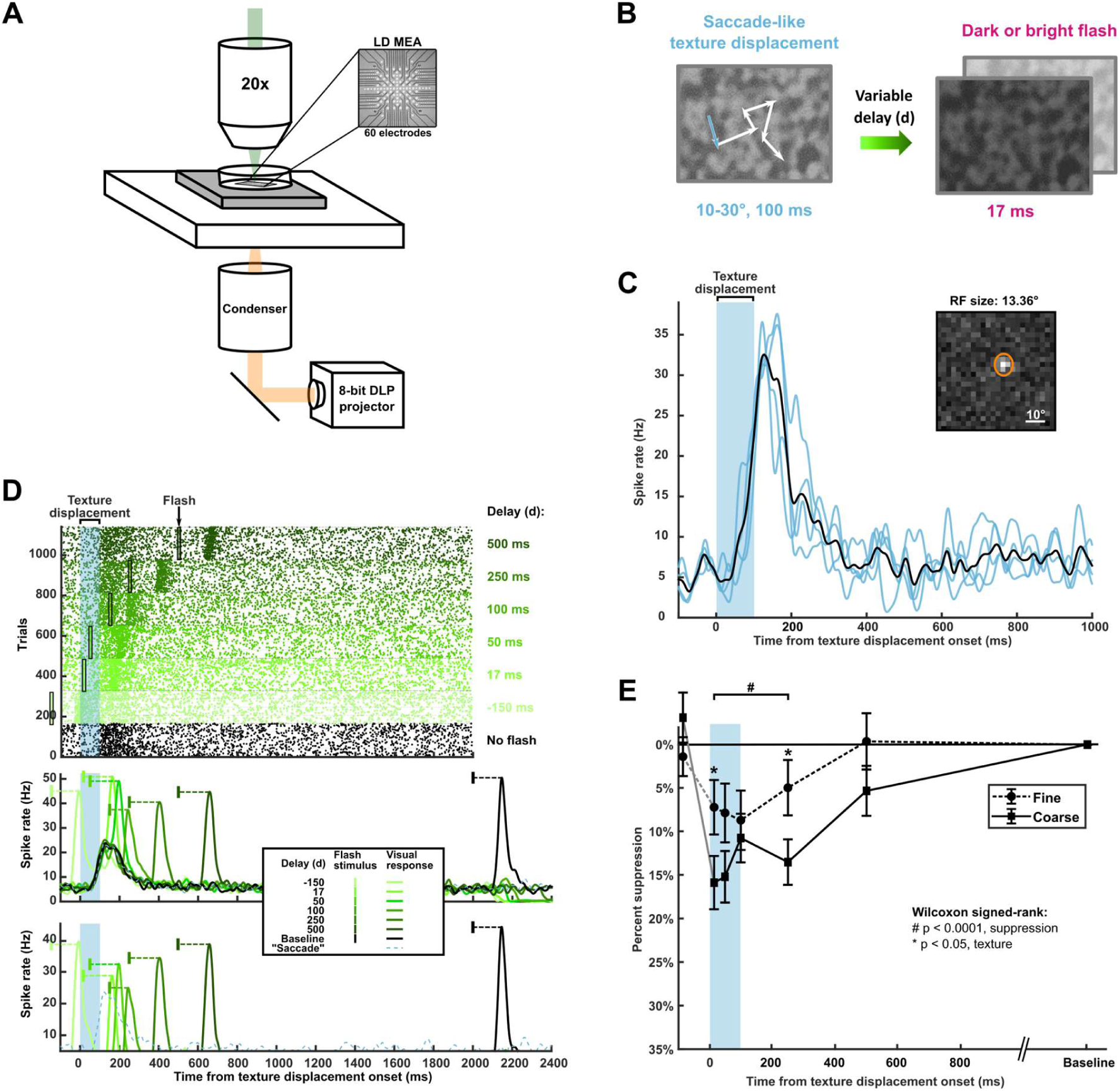
Visual saccadic suppression in zebrafish. **(A)** RGC activity of adult zebrafish retinal explants was recorded on low-density multi-electrode array (LD MEA) with simultaneous visual stimulation with DLP projector from below. (**B**) Textured stimulus was rapidly displaced in a saccade-like manner (blue arrow). After a variable delay period (d), a whole-field dark or bright flash was shown. **(C)** Spike count for individual trials (upper panel) and average spike rate across trials (middle panel) for the same example RGC as in (C), when texture displacement was paired with a whole-field flash after various delay periods. Isolated flash responses (lower panel) were acquired by subtracting “saccade” activity (blue dashed line) from flash activity (solid green lines). Green bars indicate time points of flash presentation; gray bar indicates time span of texture displacement. **(D)** Average spike rate (black line) to texture displacement of example RGC across multiple trials (blue lines). Blue bar indicates time span of texture displacement. Inset: receptive field size of that same RGC (see Fig. S1 and Methods for details). **(E)** Percent suppression (mean ± SEM) of visual responses to whole-field flash around the time of a saccade-like texture displacement (gray bar) across all RGCs recorded. RGC activity was significantly suppressed during and up to 250 ms after texture displacement onset (# p < 0.0001, two-tailed Wilcoxon signed-rank test). Suppression was significantly stronger for coarse textures, than fine (solid vs. dashed line), where indicated by asterisk (* p < 0.05, two-tailed Wilcoxon signed-rank test, Bonferroni corrected for n = 18 conditions (6 delays vs. baseline for coarse texture, 6 delays vs. baseline for fine texture, 6 coarse vs. fine texture at each delay time)). *N = 87 RGCs from 10 animals*.

**Fig. 3.**
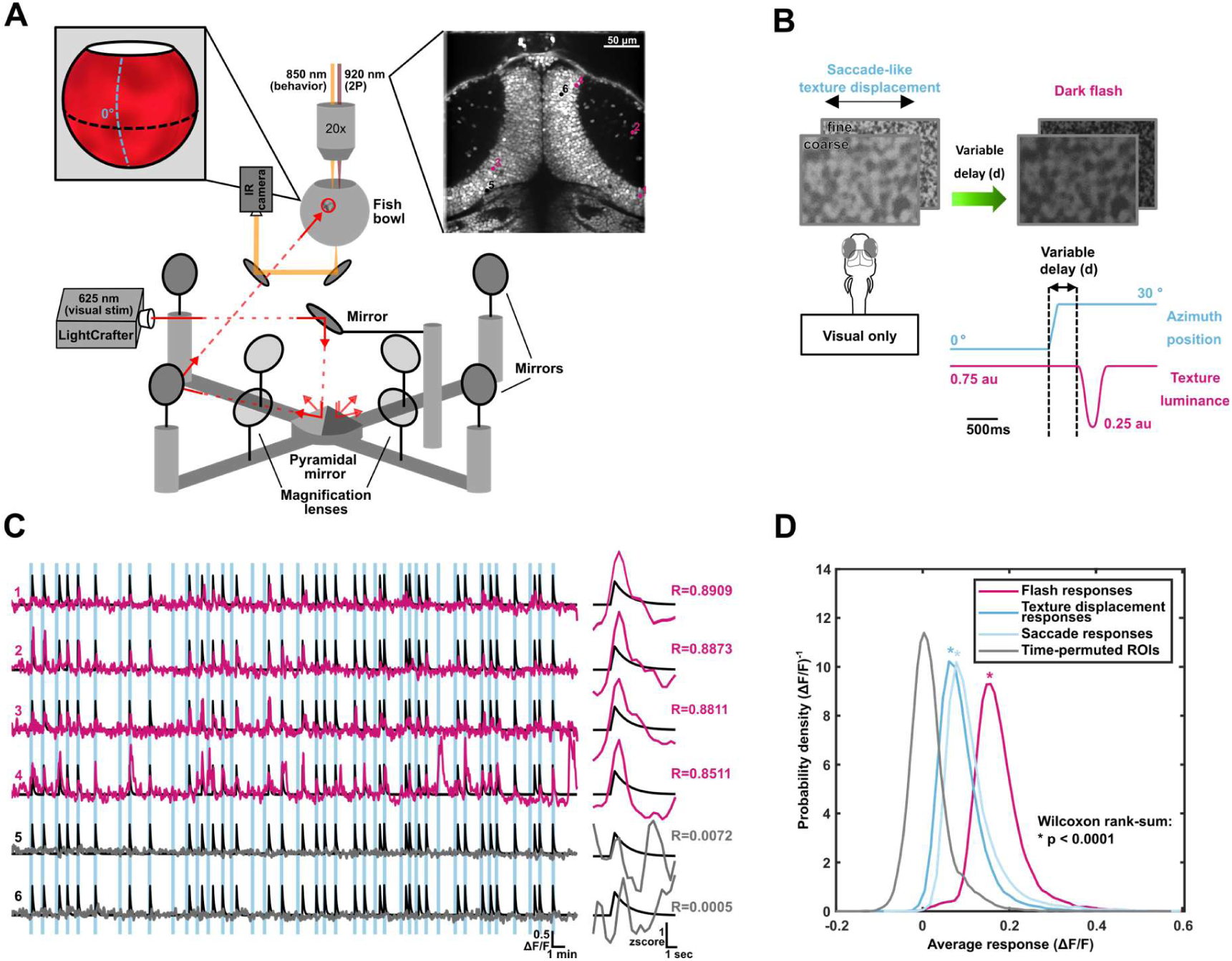
Novel visual stimulation paradigm to study saccadic suppression in zebrafish optic tectum. **(A)** Visual stimulation setup allows for whole-field visual stimulation onto a spherical projection surface, which encloses a head-restrained, live and behaving zebrafish larva (the left inset shows a photo (cropped red region) of the visual stimulus projected onto fish bowl). During visual stimulation eye position can be tracked using an infrared camera and activity of tectal neurons can be measured using 2P-imaging (right inset). Stimulus light was split by pyramidal mirror into four optical paths projecting onto the fish bowl screen surface. **(B)** Coarse and fine textures were displaced in azimuth on the spherical projection screen in a saccade-like manner. After a variable delay period (d), a dark probe flash was shown by reducing texture luminance. **(C)** Regression-based analysis of fluorescence time traces of individual ROIs (each representing one tectal neuron) revealed many neurons responding significantly to the dark probe flash, while some neurons did not respond to the dark flash. The left panel shows ΔF/F traces of example neurons from the tectal slice shown in (A), across the entire recording; the right panel shows *z*-scores of responses to a flash shown in isolation for the same ROIs. Black lines indicate the stimulus regressor, blue bars indicate the time points of texture displacements and pink and gray traces indicate neuronal activity. Pink ROIs correspond to neurons with strong correlation coefficient, R to the stimulus regressor, gray ROIs represent neurons without strongly correlated responses to the visual stimulus. **(D)** Average flash responses (pink) of all real ROIs recorded in the tectum are larger compared to average texture displacement responses (darker blue). Simulated ROIs with responses that were circularly permuted in time showed no responses (p < 0.0001 for all real responses compared to time-permuted ROIs, two-tailed Wilcoxon rank-sum test, *n = 26,556* ROIs for flash and texture displacement, *n= 14,668 ROIs* for real saccades). We also show the responses to real saccades (light blue line) here, because responses to real saccades were available from a separate dataset, which is presented in detail in Fig. 5 below.

To ensure that stimulus parameters were well matched with the RGC receptive field properties of zebrafish, we additionally characterized receptive field sizes, response latencies and response polarities of individual RGCs (Fig. S1). Adult zebrafish RGCs had receptive field sizes of 21.1 ± 6.6° (mean ± SD, n = 87) (Fig. S1A). The coarse and fine textures used in this experiment had blob sizes of approximately 14.3° and 3.6° respectively (Methods), falling within the range resolvable by RGCs. With whole-field contrast steps, the preferred polarity of RGC responses was determined (Figs. S1B, C). Most RCGs were characterized as OFF cells and preferred a bright-to-dark contrast step (72.4%, 63/87 RGCs). Fewer RGCs were characterized as ON cells (27.6%, 24/87 RGCs). For the texture displacement experiment, RGCs were only analyzed for their preferred flash polarity; ON cells for bright flashes and OFF cells for dark flashes.

In summary, we showed with visual stimuli adapted to the visual acuity of zebrafish, that zebrafish retinae modulate visual responses around the time of a saccade-like texture displacements with similar temporal dynamics as mammalian retinae. While suppression strength is reduced, compared to mammals, spatial frequency dependencies remain intact.

### Tectal neurons respond to dark flash stimuli in a 360° surround arena

The optic tectum is the major retinorecipient area in the zebrafish brain [9]. To investigate how the above-described retina-modulated responses are further processed in higher visual processing areas, we used two-photon calcium imaging to image large populations of tectal neurons in larval zebrafish. In theory, it would be ideal to compare retina and tectum at the identical developmental stages using the same activity recording technique, but due to technical limitations our comparison is limited to adult retina electrophysiology and larval tectum calcium imaging. Due to the extremely large visual field of zebrafish [31,32], it is vital to provide true whole-field visual stimulation, when studying saccadic suppression. Our recently established *in-vivo* visual stimulation setup allows for high-resolution 360° whole-field visual stimulation during calcium imaging (Fig. 3A). This enables us to investigate the consequences of saccade-like texture displacements, analog to the retinal experiments at the next step of visual processing. Tethered animals were again presented with global coarse and fine textures. Textures were displaced horizontally in a saccade-like manner and at various delay times a probe flash was shown. Probe flash parameters were slightly adjusted, to optimize for calcium imaging in the OT and closely resembled the global dark flashes used in the behavioral recordings (Fig. 3B). The duration of the flash was extended to 500 ms to allow for the relatively slow calcium signal to integrate and only dark flashes were presented, since our own observations showed that most OT neurons preferred dark stimuli over bright. Calcium recordings were registered and segmented into individual ROIs, each representing one neuron (Methods). For each ROI, ΔF/F was computed, and regression-based analysis revealed 628 ROIs (out of 26,556 ROIs, 2.36%) showing significant and reproducible responses to the flash stimulus (Fig. 3C). These responses are truly stimulus-associated, as comparison with time-permuted traces showed (Fig. 3D), thus further allowing us to investigate how these visual responses are modulated by a texture displacement.

### Texture displacement suppresses visual responses to global dark flash in the optic tectum

To investigate how texture displacements modulate visual responses in the OT, we normalized flash responses around the time of saccade to each ROI’s response to a flash shown in temporal isolation. Analogous to our analysis in zebrafish retinae, any responses associated with the texture displacement (Fig. 3D) were removed and remaining flash responses were quantified (Fig. 4). When looking at all flash-responding ROIs, we can see that more than 75% of neurons showed overall flash responses that were modulated downward, thus suppressed, by the presence of a texture displacement (Figs. 5A, S2A). We sorted cells according to their overall modulation across all flash delays and divided them into 5 bins each carrying 20% of the data. Within each bin, we sorted cells further by their most modulated flash delay in an effort to characterize the relevant delay times for which saccadic suppression or enhancement is present, and to visualize response variation across cells (Figs. 5B & S3). When comparing the 20% most suppressed and most enhanced cells to the same dataset with shuffled delay identities, we can see that, not only do responses appear more suppressed in the real data (more green coloration in Fig. 4B, left), but enhancement seems much weaker in the real data than expected by chance (less orange coloration in Fig. 5B, left).

**Fig. 4.**
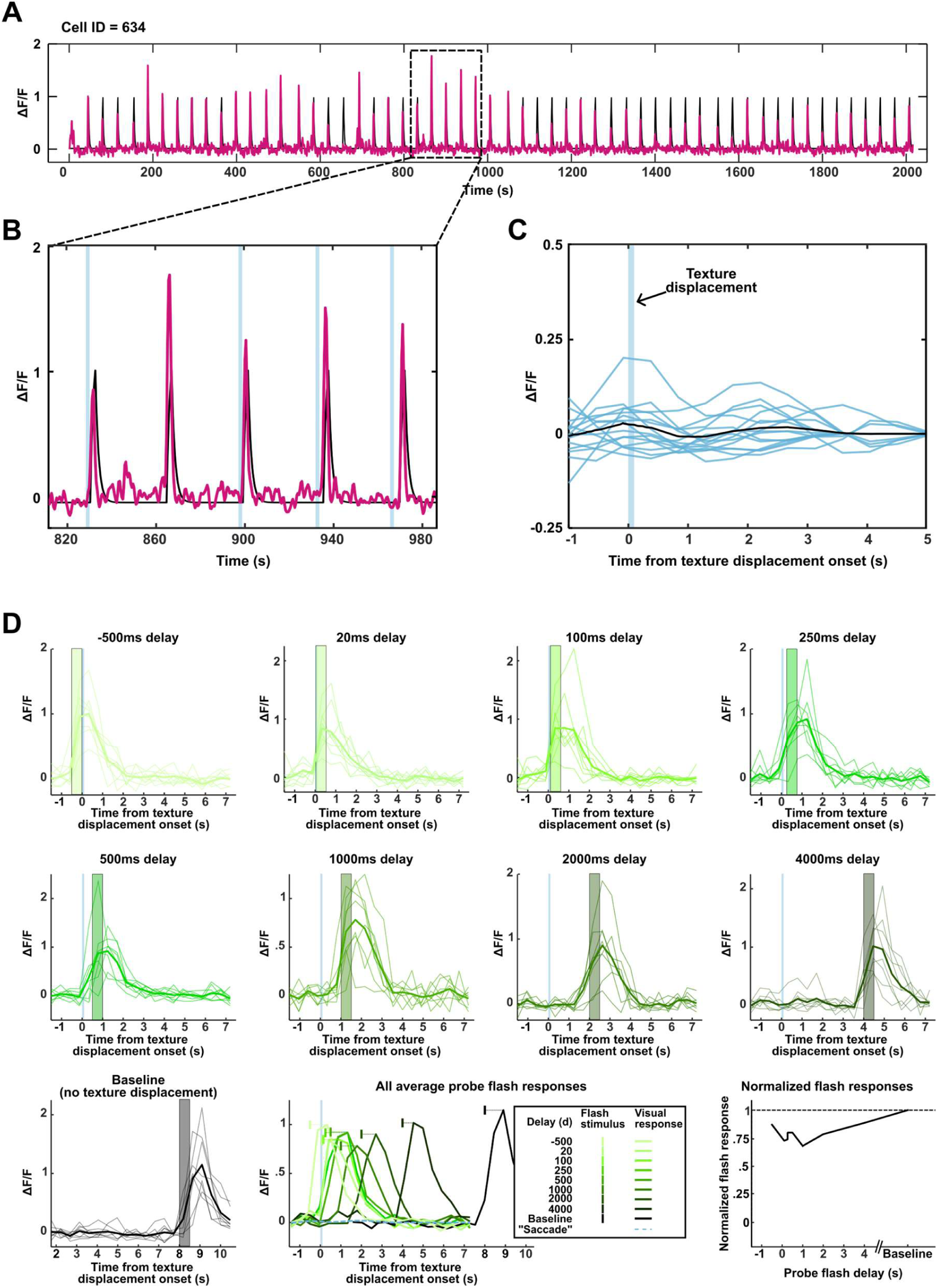
Calcium data of representative ROI. (**A**) Normalized fluorescence trace (ΔF/F) of representative ROI across an entire recording session (pink) overlayed onto the probe flash stimulus regressor (black). The ROI shows consistent stimulus-associated increases in Ca-activity of various sizes. (**B**) Zoomed in section of (**A**) with time points of texture displacement added in light blue. (**C**) Texture displacement associated responses of the same ROI shown in (**A**) and (**B**). Individual trials are shown in blue, with the average plotted in black. Texture displacement responses are much smaller compared to probe flash responses. (**D**) Calcium traces of same ROI shown above sorted by delay condition. Lines with alpha transparency value show individual trials, solid lines show averages across delay times. Light blue line indicates time point of texture displacement and green/grey box shows time point of probe flash. Lower middle panel shows all average probe flash responses for all stimulus delays and baseline condition. Lower right panel shows individual suppression curve for this ROI.

**Fig. 5.**
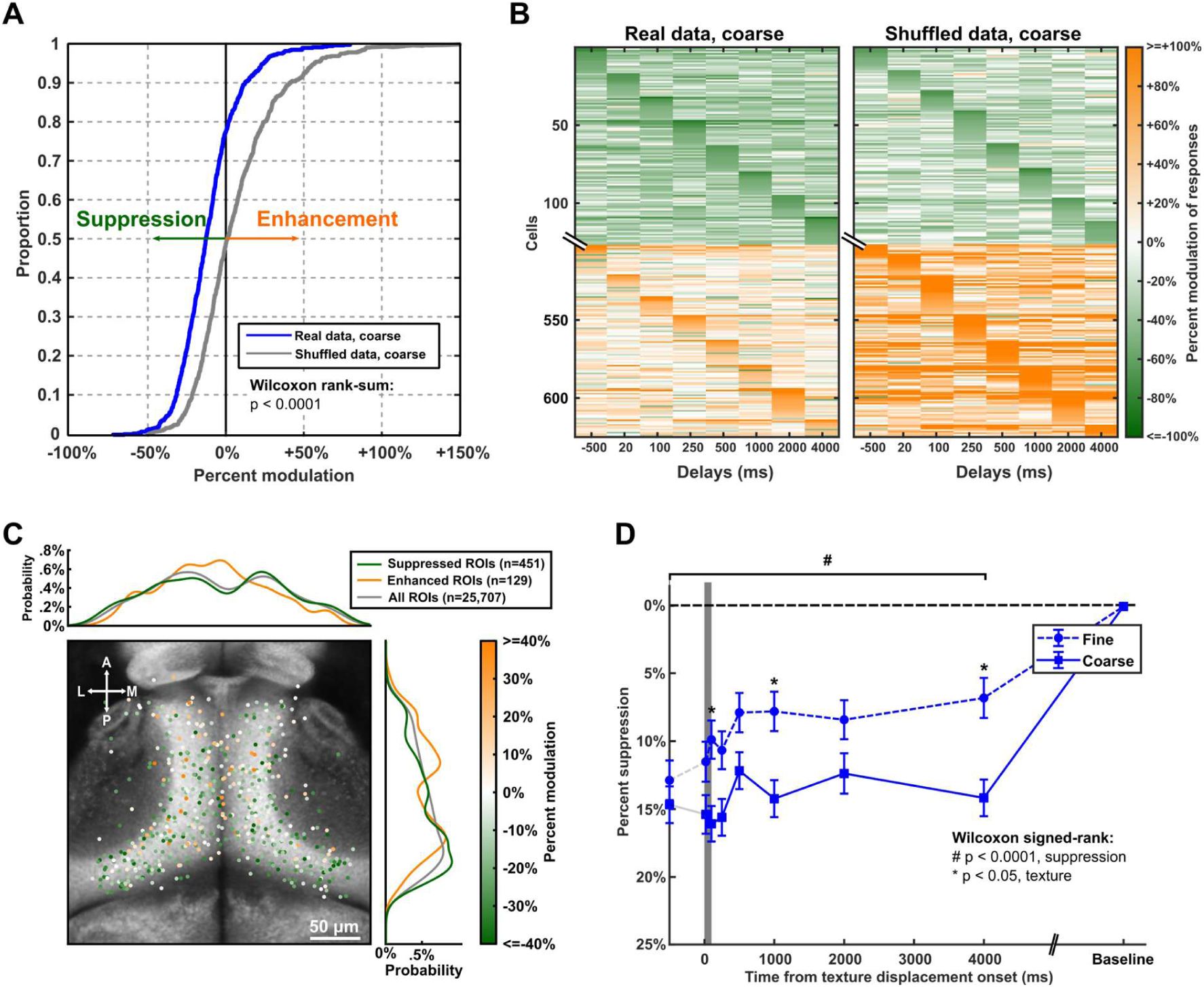
Prolonged suppression in the OT after texture displacement. **(A)** Cumulative distribution of flash response suppression across all flash-responsive neurons. Neurons were suppressed (or enhanced) by displacement of the coarse texture in a saccade-like manner (all delay times pooled). The majority of neurons have suppressed flash responses (curve for real data is shifted to the left relative to the curve for data with shuffled delay/baseline identities for individual trials, p < 0.0001, two-tailed Wilcoxon rank-sum test, *n = 628 ROIs*). **(B)** Modulation of flash responses at various flash delays of the 20% most suppressed and 20% most enhanced ROIs. Real data (left panel) shows stronger suppression and much weaker enhancement, compared to data with shuffled delay/baseline identities for individual trials (right panel). On the y-axis cells are sorted according to strength of suppression/enhancement, peak delay time, and response magnitude at peak delay (for details see Fig. S2 and methods description). **(C)** Center positions of all flash-responsive ROIs. Flash-responsive ROIs can be found all over the tectum, as is expected from the distribution of all registered ROIs in the tectum (p = 0.08 for enhanced and suppressed ROIs vs all ROIs on anterior-posterior-axis, p = 0.62 for enhanced and suppressed ROIs vs all ROIs on medio-lateral-axis, p = 0.06 for suppressed and enhanced ROIs vs all ROIs on dorso-ventral-axis (see Fig. S3), two-tailed Wilcoxon rank-sum test). Suppressed ROIs can be found more posteriorly compared to enhanced ROIs (p < 0.001 for suppressed vs enhanced ROIs on AP axis, p = 0.44 for suppressed vs enhanced on ML-axis, p = 0.99 for suppressed vs enhanced ROIs on DV-axis (see Fig. S3), two-tailed Wilcoxon rank-sum test). **(D)** Dependence of percent suppression (mean ± SEM) on delay time for all significantly suppressed neurons. Flash responses were significantly suppressed where indicated by a hash symbol (# p < 0.0001, two-tailed Wilcoxon signed-rank test, Bonferroni corrected for n = 24 conditions). Suppression was significantly stronger for coarse textures, than fine (solid vs. dashed line) for all delay times (* p < 0.05, two-tailed Wilcoxon signed-rank test). *N = 371 ROIs from 6 animals*.

We next tested whether the temporal profile of saccadic suppression is simply inherited from the retina or does change in the tectum. From Fig. 5B it is apparent that many neurons showed their strongest suppression at 1000 ms or later after texture displacement onset. This was quite surprising to us as both behavioral (Fig. 1B) and retinal (Fig. 2E) suppression usually end before 1000 ms (∼250 ms in our dataset). Texture displacements seem to modulate visual responses in the larval OT in a less straightforward fashion than in the adult retina. This can not only be seen by this very late suppression in Fig. 3B, but also when looking at the whole population of visual neurons in the OT. While there are, as previously stated, many more neurons that appear suppressed than enhanced, a spectrum of modulation exists, with many neurons also appearing enhanced or not modulated at all (Fig. S3A).

When looking at the anatomical distribution of neurons that responded to the flash stimulus, we can see that they are evenly distributed across the tectum, as is expected with a global stimulus. However, we found that neurons with suppressed responses are more likely to be found in the posterior tectum covering the temporal visual space, whereas enhanced neurons are more likely to be found in the anterior tectum, covering nasal visual space (Figs. 5C, S3).

To further investigate the temporal dynamics of saccadic modulation in the OT of not all, but only suppressed neurons, we used a permutation test to identify truly suppressed neurons independent of their actual temporal dynamics (Methods, Fig. S4). Of the flash-responsive neurons 59.1% (371/628) were significantly suppressed and these neurons do in fact appear to inherit retinal suppression for delays up to 500 ms: peak suppression for coarse and fine texture before 500 ms is nearly identical to retinal suppression (Fig. 3D, compare to Fig. 1E), with 16.1 ± 1.3% suppression for coarse and 11.5 ± 1.5% suppression for fine textures in the OT, compared to 15.8 ± 3.0% and 8.6 ± 3.4% in retina (mean ± SEM for all). The spatial frequency dependency observed in the retina remains intact as well. Unlike in retina, however, we can see strong pre-saccadic suppression in the OT. In the absence of any actual motor signal, this can only be attributed to stimulus-stimulus interaction within the tectum. Further, at delays ≥500 ms from texture displacement onset, a secondary suppression occurs, which cannot be found in the retina, with its peak at 4000 ms and 14.2 ± 1.5% (mean ± SEM) suppression of visual responses for coarse texture displacements. Initially, not expecting to see such long-lasting suppression, we had only tested delays as long as 4000 ms. But seeing that many neurons exhibited their strongest suppression at this delay, we had to ensure that we could identify a time point at which stimulus responses returned to baseline levels. Therefore, we tested in additional recordings delay times of 8, 16, and 32 s, and we found that by 8 s delays responses returned to the same strength as responses without texture displacement (Fig. S5A). Yet, we were still startled by finding this very prolonged suppression, since saccadic suppression lasting >1 second in an ecological sense, would be quite a devastating loss of visual information. To ensure that this was not an artifact of our changes to the visual stimulus between retinal and tectal recordings or a result of the slow temporal resolution of calcium imaging, we tested in a small subset of tectal neurons a short 20 ms probe flash, comparable to the stimulus used in our retinal recordings at 50 Hz imaging rate (Fig. S5B). Here, suppression lasting longer than 1000 ms remained, indicating that our result was not an artifact of the stimulus design. One possible explanation could be that our experiment did not truly assess saccadic suppression, but rather texture displacement suppression. In the paradigm of Fig. 3, the tectum only received visual information about an apparent saccade. It is, however, known in primates, that saccadic suppression is shorter with real saccades compared to visually simulated saccades [6,20]. This is why we investigated in our next steps, if the presence of a motor signal could shorten tectal suppression to values similar to other species. After all, in ecological settings, both visual and motor signals are present to shape saccadic suppression.

### Motor signals strengthen saccadic suppression, but do not abolish late suppression

Surprised by the exceptionally long lasting extra-retinal suppression we observed in flash responses of tectal neurons following a visually simulated saccade, we tested if the presence of a motor signal could shorten saccadic suppression to an ecologically reasonable length. For this purpose, we had fish perform spontaneous horizontal saccades, either in front of a textured or uniform background. Saccades were detected online and again at various delay times after the saccade onset a global dark flash was presented (Figs. 6A, B). With the same regression-based analysis described above, visually responsive ROIs were identified and analyzed for their flash responses (Fig. 6B).

**Fig. 6.**
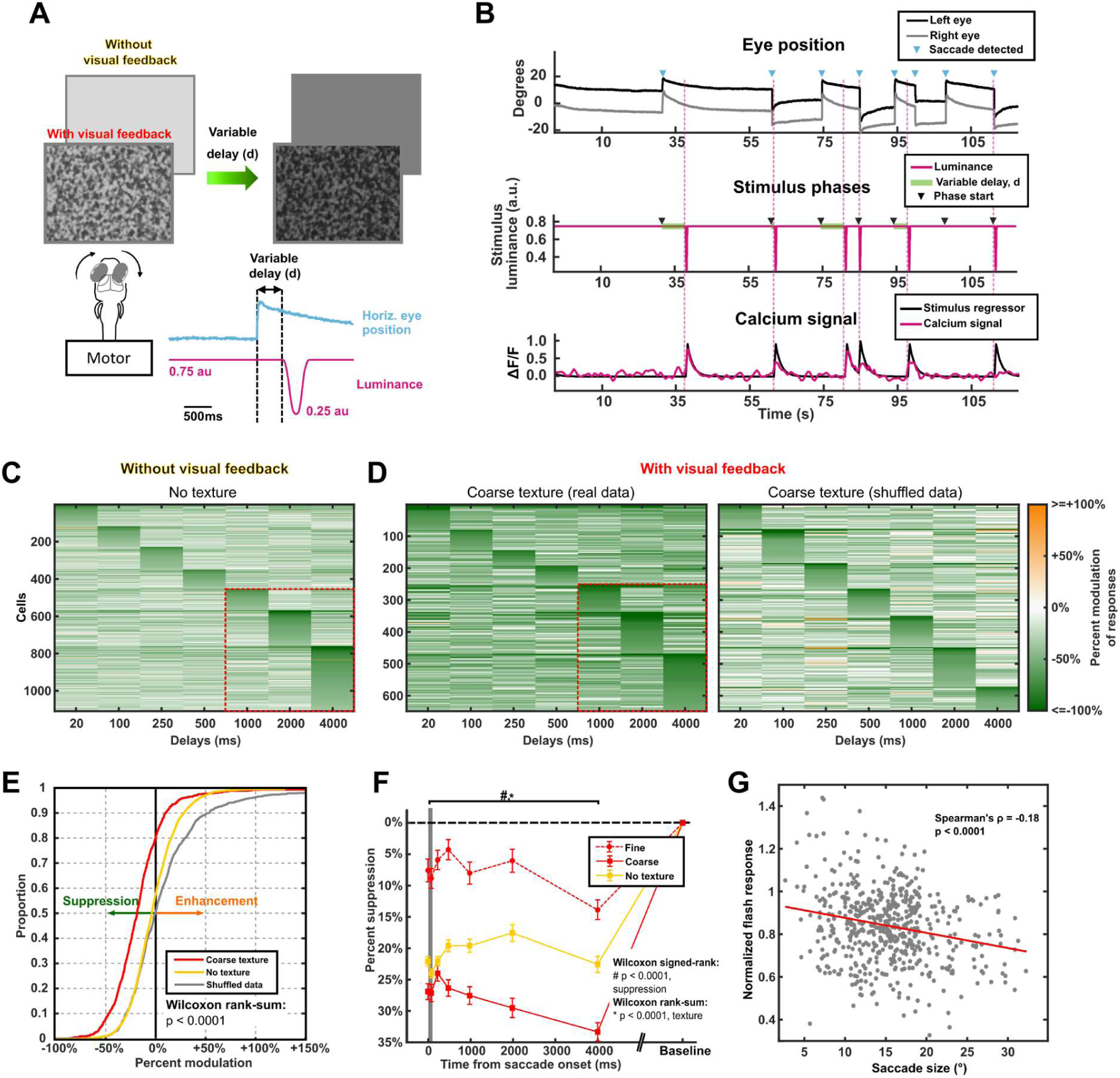
Motor signals induce strong and long-lasting tectal saccadic suppression. **(A)** Schematic of experimental paradigm. Fish performed spontaneous saccades either in front of a textured or uniform background. Saccades were detected online (blue trace) and after a variable delay period (d) a dark probe flash was shown, by reducing luminance of the texture or uniform background (pink line). **(B)** Sample recording of closed-loop stimulus design. Eye position is tracked via IR camera and saccades are detected online (upper panel). Stimulus luminance is reduced after delay period (middle panel). An example neuron responds with a weaker ΔF/F signal for shorter delay periods, than for longer delay periods (lower panel). **(C and D)** Saccade-induced modulation of flash responses at various flash delays of the 20% most suppressed ROIs, without visual feedback (uniform background, panel C) and with visual feedback (textured background, panel D left). Temporal suppression profiles were more structured than expected by chance (data with shuffled delay identities/baseline for individual trials (“shuffled data”), panel D right). Cells were sorted along the y-axis of the raster plot in the same way as in Fig. 4 and Fig. S2 (see further explanation there). **(E)** Cumulative distribution of saccade-induced modulation of flash responses in visual neurons across all flash delay times. A majority of visual neurons showed suppressed flash responses after a saccade across a coarse texture or across a uniform background; more than expected by chance (p < 0.0001, two-tailed Wilcoxon rank-sum test, n = 1,022 ROIs for textured background and shuffled data, 3,274 ROIs for uniform background). **(F)** Temporal profiles of percent suppression of response (mean ± SEM) for all neurons significantly suppressed by a saccade across coarse, fine or uniform background. Flash responses were significantly suppressed for all delay times and all backgrounds (# p < 0.0001, two-tailed Wilcoxon signed-rank test, Bonferroni corrected for n = 21 conditions). Suppression was significantly stronger for coarse than fine textures (solid vs. dashed red line) for all delay times (* p < 0.001, two-tailed Wilcoxon signed-rank test). *N = 494 ROIs from 6 animals for textured background, n = 643 ROIs from 6 animals for uniform background.* **(G)** Normalized flash response for delay times smaller than 500 ms and saccade size are negatively correlated, where larger saccades lead to more strongly suppressed flash responses (p < 0.001, Spearman rank correlation, n = 543 saccades).

Once again, visual responses are diversely modulated up or down (Fig. S2B). When looking at the 20% most suppressed neurons, it can be seen that the temporal suppression profile is more structured than would be expected by chance (compare Figs. 6C and D, left with Fig. 6D, right), with and without visual feedback from a textured background. Peculiarly, there seem to be even more neurons with strong suppression for delays ≥ 1000 ms in this condition (see red box in Figs. 6C & 6D). Consistent with our findings in simulated saccades, more neurons are suppressed than enhanced (Fig. S2B). With textured visual background, similar to the texture displacement experiment, nearly 80% of neurons appeared to be suppressed across all delay times. Without visual feedback background texture only ∼60% of neurons were modulated downwards by a saccade (Fig. 6E). Saccadic enhancement appeared to be present in some neurons as well, but enhancement was less prominent than would be expected by chance. Therefore, suppression in the OT can also be mediated by a real saccade in the absence of a structured visual surround.

The visually induced saccadic suppression, which can be seen in texture displacement experiments in retina and OT is further augmented in the presence of a motor signal. Peak suppression for early delays <500 ms is increased by nearly 70% in presence of a motor signal (16.1 ± 1.3% suppression for texture displacement vs. 27.1 ± 1.4% suppression saccade (mean ± SEM for both, coarse stimulus condition)). Further, we had anticipated that through the presence of a motor signal, long-lasting suppression after >500 ms would be weakened or even abolished. As already indicated by the whole neuron population (Figs. 6C, 6D, S2B), when looking at significantly suppressed neurons only, we observed strong suppression multiple seconds after saccade onset, particularly in the textured condition, (Fig. 6F). This long-lasting suppression was present both for conditions with uniform and textured background. This experiment with real saccades showed that saccadic suppression in the OT can be accomplished by motor signals alone in absence of visual feedback. Motor-related suppression is stronger than visual suppression alone, suggesting a strong influence of a corollary discharge signal in saccadic suppression. If the purpose of such a corollary discharge signal is to act as a suppressive command for visual neurons during rapid eye movements, larger saccades, that lead to larger visual transient, should induce stronger suppression.

Therefore, we tested if motor driven saccadic suppression in the OT might also correlate with saccade size. Particularly for early delay times <500 ms, saccade size is indeed negatively correlated with normalized flash response, thus for greater saccade sizes, responses are down-modulated more strongly or suppressed (Fig. 6G), not however for late delay times ≥500 ms (Fig. S6). This is in congruence with recent findings of a corollary discharge signal of saccade-like swim bouts in the zebrafish OT, which in size is positively correlated with swim vigor [21].

In summary, saccadic suppression in the OT can be elicited by visual or motor effects alone, and it is strongest in the presence of a motor signal. Suppression strength is correlated with saccade size, indicating that a corollary discharge signal is graded to the expected sensory signal that is to be suppressed. And lastly, both simulated and real saccades induce long lasting (>500 ms) suppression for global flash stimuli.

### Saliency filtering can explain long lasting suppression in OT

After having found that motor signals can enhance early saccadic suppression but cannot abolish long-lasting suppression to a global flash stimulus, we considered the behavioral context of saccades in larval zebrafish. Larval zebrafish will perform non-reflexive saccades primarily to reorient their gaze and examine different parts of the visual field or in the context of prey capture to locate prey objects on the area centralis [15,33]. In both cases, it is beneficial for the animal to focus visual attention onto local stimuli, such as an approaching predator or a small prey object. Now, if the animal was visually inhibited, due to saccadic suppression lasting several seconds past an actual saccade, behaviorally highly salient stimuli like predator and prey, could not be detected. Therefore, we suspected that late suppression, lasting >1000 ms could act like a visual saliency filter, ensuring efficient sensory processing, rather than saccadic suppression per se. While behaviorally relevant local stimuli could remain fully visible, less relevant global stimuli would be suppressed.

To test this hypothesis, we combined a texture displacement with a local prey-like moving dot (Fig. 7A) or a looming disc, simulating an approaching predator (Fig. 7B), rather than a global flash, as in previous experiments). Global flashes do occur frequently in water bodies due caustics and refraction of light at the water surface [34]. This flickering should mostly be ignored by the animal, since it is non-informative (although we saw in Fig. 1 that sparse flashing will induce O-bend behavior). Due to association of moving dot stimuli with hunting behavior, and looming stimuli with escape behavior, these two stimuli are arguably more ecologically relevant than global flash stimuli. The local dot stimulus elicited visual responses in more anterior-medially localized neurons, as is expected, due to the tectal retinotopy [35] (Fig. S7A, corresponding to stimulus positions in the upper-nasal visual field), while the looming disc stimulus elicited responses more ventrally (Fig S7B, corresponding to lower elevations). Moving dot responses were comparable in size to global flash stimuli in the previous experiments, while looming discs evoked larger responses compared to both (Fig. S8). As hypothesized, visual responses to the ecologically relevant stimuli were barely modulated across all delay times. We did not see more suppressed than enhanced neurons, contrasting the suppression profile observed in our previous global flash experiments (Fig. 7C). When looking at the significantly suppressed neurons only, we can see that for both behaviorally relevant stimuli there is only weak suppression for early delay times; late suppression is completely abolished (Fig. 7D). This early suppression is likely inherited directly from the retina and it follows a roughly similar time course and suppression strength as observed in our adult retinal recordings (compare black lines in Fig. 8A).

**Fig. 7.**
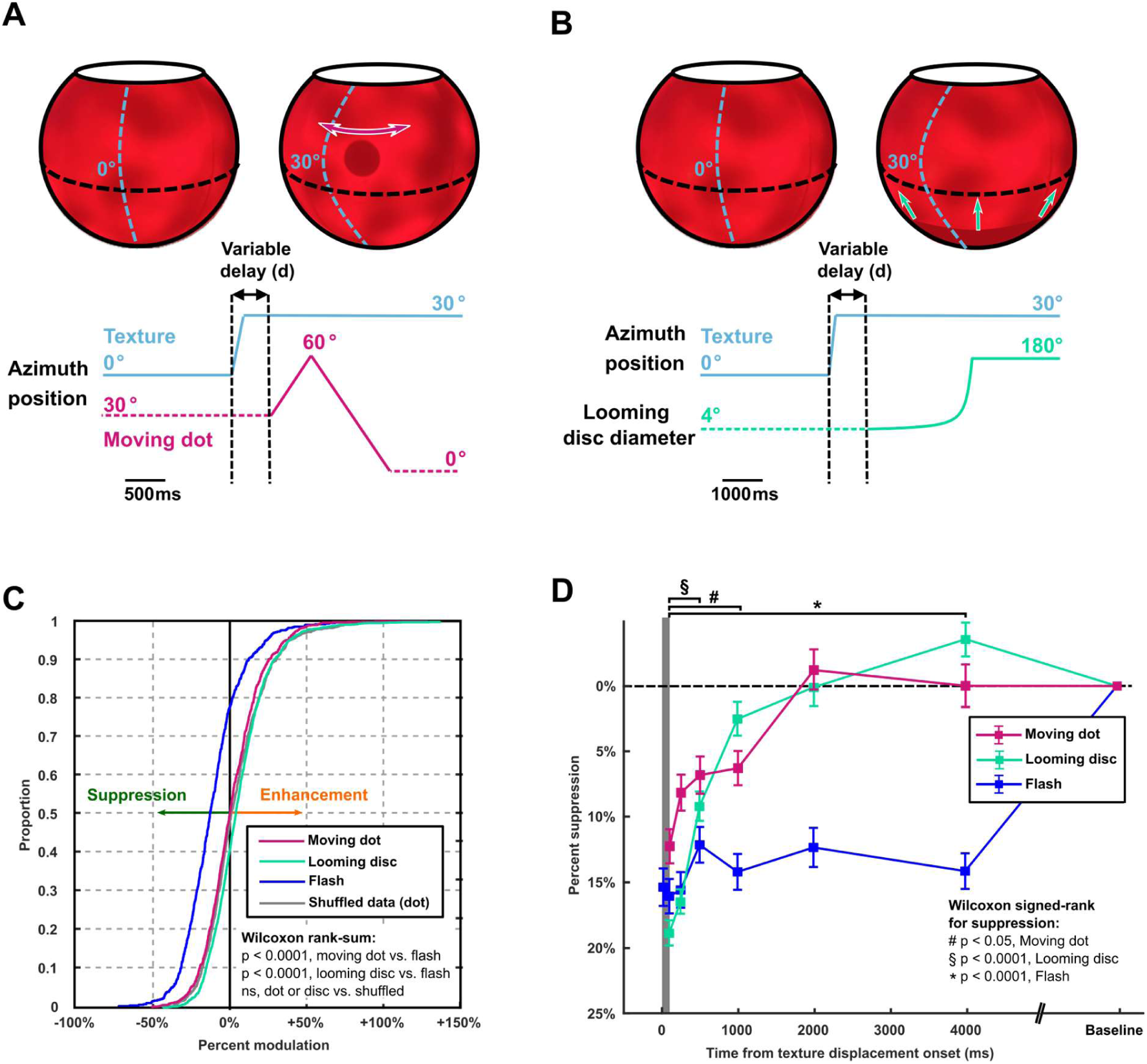
Responses to local stimuli persist after simulated saccades. (**A**) Texture displacement was paired with a local prey-like moving dot stimulus as a probe stimulus. Coarse texture was displaced in a saccade-like manner (blue line) and after a variable delay period (d) a dark dot with a diameter of 20° was shown at 10° elevation, moving in azimuth at 60°/s across the left frontal visual field (pink line, Methods). (**B**) Texture displacement of a coarse texture (blue line) was paired after a variable delay with an expanding disc stimulus, simulating and approaching predator. Disc diameter expanded from 4° to 180° originating from the south pole dark shade and green arrows (Methods). (**C**) Cumulative distribution of texture displacement-induced modulation of moving dot and looming disc responses in visual neurons across all flash delay times. Unlike in experiments with a global flash probe stimulus (blue line, from Fig. 3A), for the moving dot stimulus (pink line) and the looming disc stimulus (green line), visual neurons showed roughly equal amounts of suppressed and enhanced responses following texture displacement as expected by chance (p < 0.0001 for moving dot vs. global flash, p < 0.0001 for looming disc vs. global flash, p > 0.05 for moving dot vs. data with shuffled delay identities for individual trials (“shuffled data”) and looming disc vs. shuffled data, two-tailed Wilcoxon rank-sum test, *n = 956 ROIs* for dot*, n = 1,827 ROIs* for looming disc). (D) Temporal profile of response suppression for moving dots (pink) and looming disc (green) (mean ± SEM) for all neurons suppressed by a displacement of a coarse texture. Moving dot responses are only suppressed for delay times ≤1000ms, looming disc responses are suppressed for delays ≤500ms (# p < 0.05 for moving dot, p < 0.0001 for looming disc, two-tailed Wilcoxon signed-rank test, Bonferroni corrected for n = 18 conditions). Suppression for dot stimulus is weaker than for flash and long-lasting suppression is abolished (* p < 0.0001 for dot vs flash, p > 0.05 for dot suppression 2000 ms and 4000 ms, two-tailed Wilcoxon rank-sum test). Suppression for looming disc for early delays is as strong as flash suppression, but long-lasting suppression is abolished (p > 0.05 for disc vs. flash for delays ≤500ms, two-tailed Wilcoxon rank-sum test). *N = 312* ROIs from 3 animals for moving dot, *n = 145 ROIs* from 5 animals for looming disc.

**Fig. 8.**
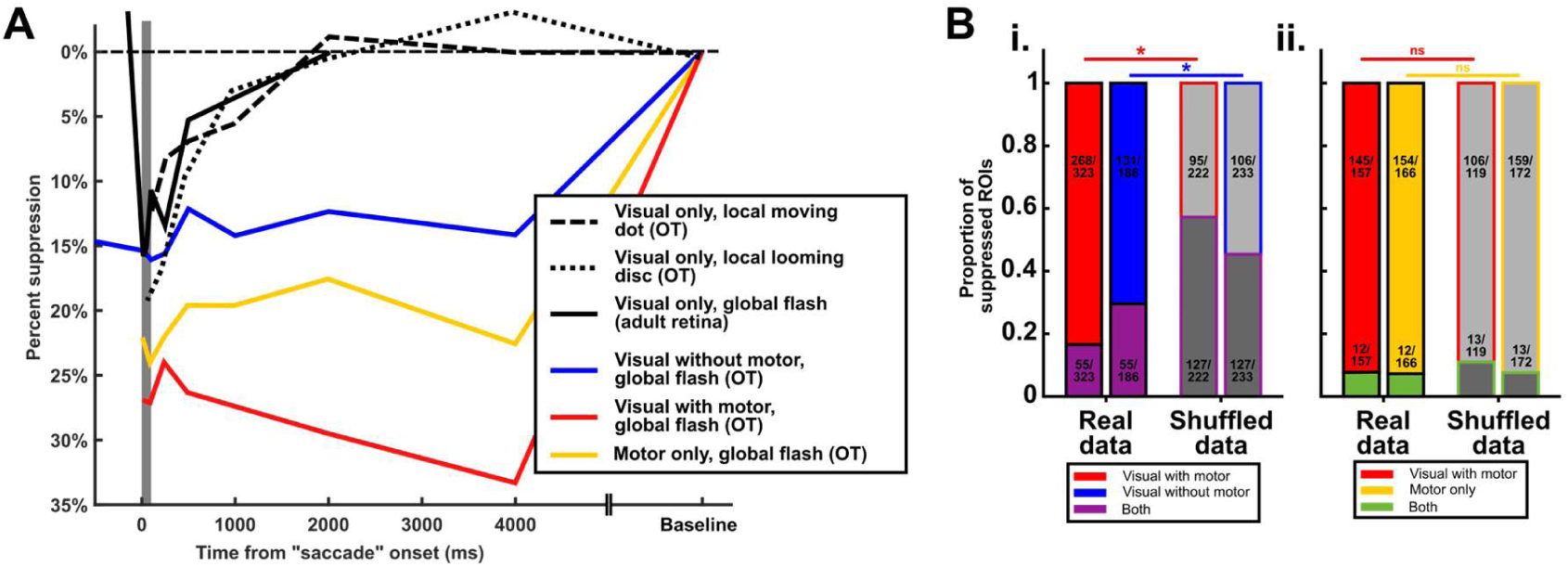
Visual and motor signals act on non-overlapping neural populations in the OT. **(A)** Temporal suppression profiles in retina and OT of all visual and motor conditions tested in this study. Visual Only refers to experiments with simulated saccades on a textured background, Visual & Motor Combined refers to experiments with real saccades across a textured background, and Motor Only refers to experiments with real saccades across a uniform background. **(B)** i: Proportions of neurons suppressed exclusively in the Visual & Motor Combined condition (red) or exclusively in the Visual Only condition (blue) versus neurons suppressed in both conditions (purple) when tested within the same animal, compared to the proportions in data with shuffled condition identities. There are less Visual & Motor, exclusive suppressed neurons than expected by chance (p < 0.0001, χ^2^-test for proportions) and less Visual Only, exclusive suppressed neurons than expected by chance (p < 0.0001, χ^2^-test for proportions). ii: Proportions of neurons suppressed exclusively in the Visual & Motor Combined condition (red) or exclusively in the Motor Only condition (yellow) versus neurons suppressed in both conditions (green) when tested within the same animal, compared to the proportions in data with shuffled condition identities. Only few neurons are suppressed in both conditions, no more than expected by chance (p = 0.91, χ^2^-test for proportions), suggesting that they are separate populations.

Overall, we show that late saccadic suppression lasting >1000 ms is heavily dependent on the probe stimulus used. While global stimuli, with little to no saliency for the animal, remain suppressed for several seconds after a saccade, behaviorally more relevant local stimuli, such as a prey-like dot or a looming disc simulating a predator, become quickly perceivable again. This task-specific processing of visual information in the tectum could function as a saliency filter, ensuring efficient sensory encoding for zebrafish that have limited neural capacities.

### Saccadic suppression is shaped by motor signals and the type of visual stimuli in zebrafish OT

When combining the observations from all experiments described above, we can identify both visual and motor-related mechanisms that shape saccadic suppression dynamics across the zebrafish visual system (Figs. 8A, 9). In isolated adult retinae, RGC responses to global flash stimuli are weakly suppressed for up to 250 ms; the suppressed signal is relayed to the OT. Here, visual responses of larval tectal neurons are even more strongly suppressed by a real saccade, than a simulated saccade. At later time points (≥1000 ms) after saccade onset, high and low saliency visual stimuli are differentially processed by tectal neurons. While responses to behaviorally less relevant global flashes are strongly suppressed for several seconds, responses to a local prey-like dot or a looming predator, after weak early suppression in the first 1000 ms after saccade onset, quickly return to baseline response level. Lastly, when investigating motor and visual effects in neurons of the same animal, we found non-overlapping populations of neurons. Only a small fraction of neurons (55/323 ROIs and 55/186 ROIs) were suppressed in both “visual without motor” and “visual with motor” conditions, less than expected by chance. Instead, most neurons were either exclusively suppressed in the presence of a motor signal (268/323 ROIs, 83.0%, Fig. 8Bi, “Visual with motor”) or in the absence of a motor signal (131/186 ROIs, 70.4%, Fig. 8Bi, “Visual without motor”). When comparing saccades with visual feedback to saccades without visual feedback in neurons of the same animal, we again found that almost all neurons were exclusively suppressed in either one of the two cases, saccades with visual feedback (145/157 ROIs, 92.3%, Fig. 8Bii, “Visual with motor”) or saccades without visual feedback (154/166 ROIs, 92.8%, Fig.86Bii, “Motor only”).

**Fig. 9.**
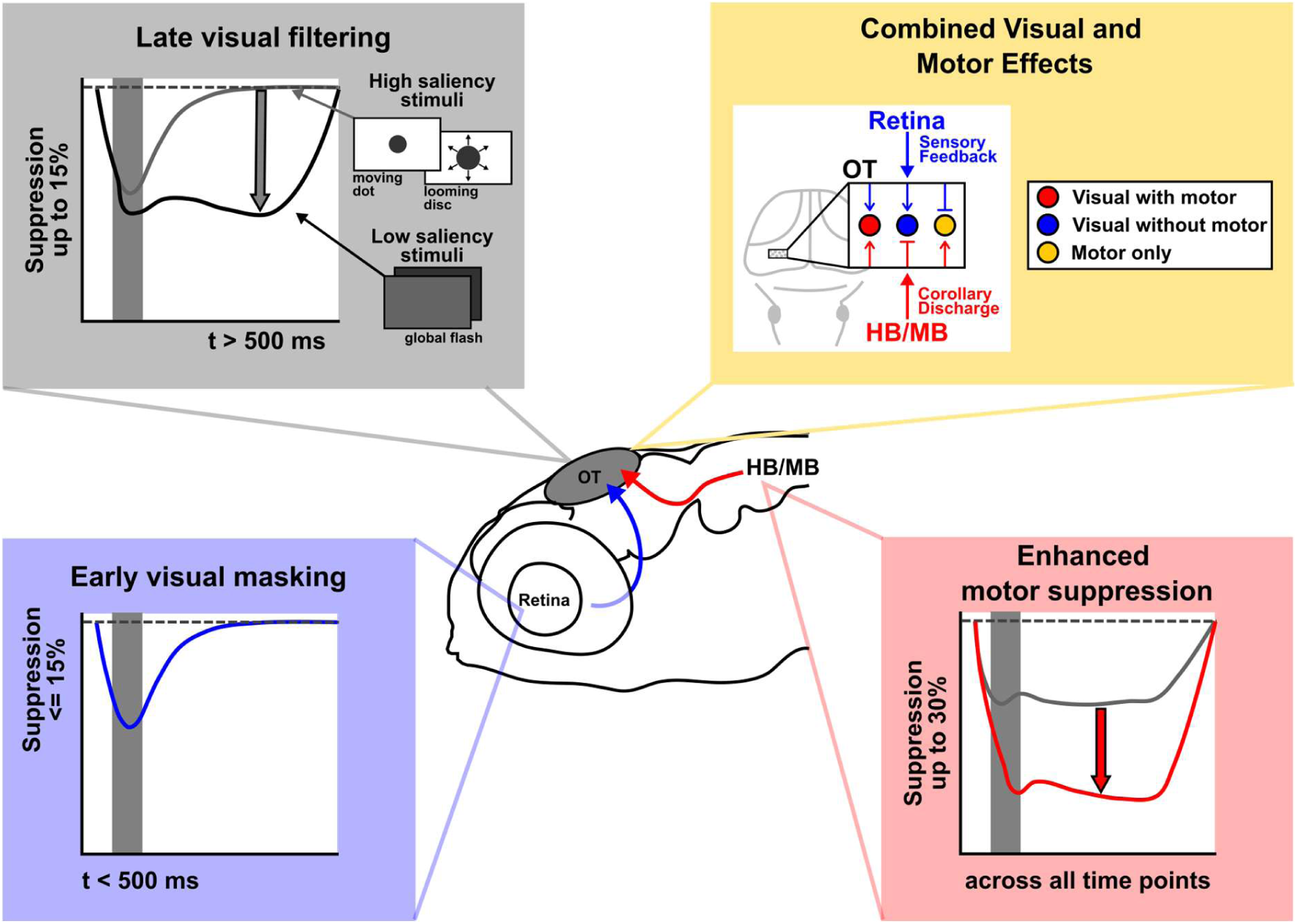
Visual and motor mechanisms shape saccadic suppression throughout the zebrafish visual system. Both visual and motor-related signals shape the temporal profile of saccadic suppression in the zebrafish. Purely visual mechanisms (i.e. visual masking) lead to early, weak saccadic suppression in the retina (blue box). Retinal signals are sent to the OT and further modulated. Motor signals, likely originating from the hindbrain oculomotor circuits, strengthen saccadic suppression in the OT (red box). Visual filtering mechanism in the OT can alter late saccadic suppression to last more than 1000 ms (gray box). Behaviorally salient stimuli, such as a moving prey-like dot or a looming disc, simulating an approaching predator, are not further suppressed, whereas low saliency stimuli, such as a global flash are suppressed for long time scales (gray box). Saccadic suppression of separate neuronal populations is context-specific. Some neurons receive sensory input, some motor input and others require both to be saccadically suppressed (yellow box). Overall tectal suppression is strongest when sensory and motor inputs are combined.

Very few neurons were suppressed for both conditions, no more than expected by chance (Fig. 8B., ii.). These findings suggest that saccadic suppression manifests in separate neuronal populations depending on visual and motor context.

## Discussion

Motivated by having found evidence for behavioral saccadic suppression in larval zebrafish, we identified distinct visual and motor-related processes, shaping saccadic suppression on a neuronal level across the zebrafish visual system (Fig. 9). Purely visual effects, such as masking, already take place in zebrafish retina prior to entering the OT. Following a saccade-like texture displacement, RGCs show suppressed responses to peri-saccadic visual stimuli. This effect is not specific to texture displacements. In mammalian retinae and human psychophysics experiments, simple luminance steps are known to suppress responses to probe flashes within a similar time window as saccadic suppression [6]. We found that pairing two probe flashes can lead to very weak suppression of tectal responses to the second probe flash, lasting up to 500 ms, likely due to simple visual masking (Fig. S5C), similar to the masking effect seen for the local dot and the looming escape visual stimuli in the tectum.

The length of the peri-saccadic suppression window in zebrafish retina is comparable to that found in mammals, but suppression strength is much weaker [6]. Spatial frequency dependencies of suppression are similar to those observed in mammals in that low spatial frequency content is more strongly suppressed than high spatial frequency content. Zebrafish have lower visual acuity though, so the stimuli were coarser in this study than in research on mammals.

This modulated retinal signal gets further relayed to the OT, the major retinorecipient area in the zebrafish brain. There, saccadic suppression is further shaped by both visual and motor signals. Saccade-like texture displacements elicit saccadic suppression of visual responses in the OT just as in the retina.

In fact, suppression in the early peri-saccadic window (delays <500 ms) is likely directly inherited from the retina, as it follows a similar time course and has nearly identical suppression strength. When comparing purely visually induced “saccadic” suppression to saccadic suppression following an actual spontaneous saccade, we found that the execution of an actual motor command significantly increases suppression strength. This result suggests that corollary discharge signals play an important role in strengthening saccadic suppression. The observed saccade-size dependency (Fig. 6G) supports this idea. Corollary discharge signals are known to play a vital role in optimizing sensory processing during self-motion across the animal kingdom [36–38] and recently, it has been shown that the zebrafish OT receives corollary discharge signals during saccade-like swim bouts [21]. It is likely that eye movement signals could elicit a similar corollary discharge signal in zebrafish, and our observed “motor only” saccadic suppression with a homogeneous visual background suggests that these corollary discharge signals exist. Future studies are needed to measure the saccadic corollary discharge signal more directly in the zebrafish OT and demonstrate its mechanistic involvement in saccadic suppression.

On a behavioral level, in primates, motor signals are known to shorten the peri-saccadic suppression window[6], potentially due to post-saccadic excitability enhancement in cortical areas [39–41]. The lack of a cortex is thus a potential proximal explanation for the lack of a motor-associated temporal restriction of saccadic suppression in zebrafish.

Another striking difference between saccadic suppression in the zebrafish OT and saccadic suppression in primates is a strong secondary suppression in zebrafish, particularly of global flash responses for several seconds past the saccade onset. This was true for simulated saccades and real saccades with and without visual feedback. We were struck by this finding, as from an ecological perspective, it seems like a massive loss of visual information, to discard sensory signals for up to 4 seconds after a saccade. Using an imaging technique with very slow temporal resolution of ∼2 Hz, attempting to resolve an effect that ought to play in the tens and hundreds of millisecond time scale, we were concerned that the slow suppression dynamics were an artifact of our stimulus design and poor temporal resolution. However, responses to the dot or looming stimulus (with identical imaging settings) had much shorter suppression durations (Fig. 7, Fig. 8A), suggesting that the observed long delays aren’t caused by experimental artifacts. We also performed a control experiment with very brief global flashes and faster imaging, in which suppression lasting > 1000 ms remained (Fig. S5). The observed longer-lasting suppression in zebrafish could potentially be related to the much lower saccade frequencies in zebrafish vs. primates, and further studies are needed to investigate the role of suppression duration across species.

We propose that the persistent suppression is a task-specific efficient encoding strategy for the larval zebrafish, which has to deal with limited neuronal resources. Since spontaneous saccades are often followed by the detection of a local visual stimulus, either in a prey capture setting or when detecting a predator [15,33], it would be sensible for the fish to focus its neural resources to accurately detecting local behaviorally salient visual stimuli and discarding information about global stimuli. This is exactly what we found when testing a local prey-like dot stimulus or a looming disc stimulus, simulating an approaching predator for saccadic suppression in the OT. While the visual suppression inherited from the retina remains intact, there is no further modulation of the visual signal after the initial peri-saccadic window. Such attentional filtering for behaviorally salient stimuli following a saccade has already been predicted in computational models of cortical population activity; in a competitive interaction between two stimuli, modeling studies predict suppression of unattended stimuli to enhance sensitivity to relevant stimuli [42]. Further, studies in archerfish have shown that the teleost optic tectum is capable of using visually driven long-range inhibition, i.e. inhibition modulated by signals from outside the receptive field, to aid in selecting behaviorally salient stimuli, such as prey over behaviorally irrelevant stimuli [43]. In the archerfish retina, saccades might serve to encode visual stimuli with higher fidelity [44]. Overall, future studies are needed to reveal the differential suppression of various ecologically relevant visual stimuli in the context of saccadic suppression. In the cortical area V1 of primates, preliminary data suggests that dark contrasts are not suppressed at all, unlike in superior colliculus and primate perception [45,46], which could represent an ecological adaptation as well, and this phenomenon needs to be addressed in comparative studies.

Ethologically relevant cues oftentimes have higher spatial frequency content than less relevant cues (e.g. prey stimuli). Anatomical regionalization has been described for tectal neurons representing visual stimuli of different scale (ref Foerster et al.). Since stimuli of lower spatial frequency (“coarse stimulus” here) and the global flashes are associated with higher levels of saccadic suppression, this anatomical regionalization could provide a mechanism for the stimulus specificity, since suppressive inputs could be regionalized as well. Further work is needed to test this and also improve our understanding of the receptive field properties and functional roles of tectal neurons.

Lastly, we investigated how visual and motor signals can act on tectal neurons within the same animal. We have found that while there are many neurons that can be suppressed purely by a visual signal of a simulated or real saccade across a texture, the occurrence of stronger suppression requires a simultaneous motor signal, arising from an actual saccade. Further, when comparing which neurons are suppressed through the execution of a saccade in the absence of any visual feedback to those suppressed through a saccade with visual feedback, we found that there is almost no overlap between these two cell populations. It is unclear why motor suppression and visually driven suppression are segregated in such a manner. It seems that saccadic suppression is used as a precise tool for various different contexts. This suppression layout across the neuronal population could be the results of balancing suppression against information loss, since only small populations are suppressed, and only under certain conditions.

In summary, we identified that saccadic suppression exists in the small-brained larval zebrafish, and that visual and motor signals play differential roles. Compared to mammals, zebrafish saccadic suppression is weaker but longer-lasting. Further studies are needed to reveal conserved mechanisms that govern saccadic suppression and potentially explain the observed differential expression of suppression dynamics across species.

## Methods

### Zebrafish

For retinal recordings we used retinae of *nacre +/-* adult zebrafish (*Danio rerio*), which are functionally wild type. Ten retinae from 6 male and 4 female fish (age 8-23 months) were used. Adult zebrafish were housed at our local animal facility at 28°C water temperature at a 14/10h light/dark cycle. Animals were brought to the laboratory space and dark-adapted for several hours prior to retinal tissue preparation.

For behavioral recordings we used *wt* larval zebrafish, age 7 dpf. Zebrafish were reared in E3 solution at 28°C on a 14/10 light/dark cycle. Animals used for free-swimming experiments were fed once daily at 5 and 6 dpf with artemia powder (JBL GmbH, Neuhofen, Germany).

For *in-vivo* calcium recordings, larval zebrafish, age 5-7 dpf, with pan-neuronal GCaMP6f expression and homozygous for the *mitfa* mutation were used (*Tg(HuC:H2B-GCaMP6f)jf7; nacre -/-*) [49]. At 4 dpf, animals with the strongest GFP expression were selected for subsequent recordings at 5-7 dpf. Animal experiments and all experimental protocols were approved by the responsible ethics committee of the Regierungspräsidium Tübingen in accordance with German federal law and Baden-Württemberg state law. All animal procedures also conformed to the institutional guidelines of the University of Tübingen.

### Retinal electrophysiology

#### Tissue preparation (MEA)

Fish were dark adapted for a minimum of 2 hours, before they were killed under dim red light either with anesthetic overdose (250 mg/L Tricaine, Pharmaq, in fish water) or through rapid chilling (∼2-4°C). Afterwards, animals were decapitated and the eyes were removed and placed into chilled Ringer solution (in mM: 110 NaCl, 2.5 KCl, 1.0 CaCl2, 1.6 MgCl2, 10 D-Glucose, and 2.2 NaHCO3) bubbled with 5% CO_2_ and 95% O_2_. Retina was isolated inside a Ringer bath, placed into 5% Hyaluronidase solution for 3 min to remove remaining vitreous and then placed into warmed Ringer solution (∼30°C) for one hour to recover tissue. Following recovery, retina was mounted onto a nitrocellulose filter (Millipore) with a central 1.5 x 1.5 mm hole and placed RGC side down onto a perforated low-density multi-electrode array (60pMEA200/30iR-Ti-gr, Multichannel Systems, Reutlingen, Germany) with 60 electrodes in a square-grid arrangement and 200 µm inter-electrode distance. Good electrode contact was ensured through negative pressure through the perforation over at least 30 minutes, prior to recording begin. Tissue was superfused with bubbled Ringer solution at ∼30-32°C. Data were acquired at 25 kHz using MC Rack version 4.6.2 (Multichannel Systems).

#### Visual stimulation (MEA)

Visual stimuli were presented on a DLP projector with a refresh rate of 60 Hz. Visual stimuli were adapted from Idrees and colleagues [6]. A spectral intensity profile (in μW cm^-2^ nm^-1^) of the visual stimulus was measured with a calibrated USB2000+ spectrometer (OceanOptics, Orlando, FL, USA) and converted into biological units (R* rod^-1^ s^-1^) [49] normalized zebrafish rod absorbance[47](2.4 μm^2^ for zebrafish rods[48]). On the DLP projector scale (0-255) a stimulus intensity of “60” with ND5 filter (which was used in almost all retinal recordings) equaled 3442 R* rod^-1^ s^-1^.

Two binary Gaussian blur textures with differing spatial scales were used in the experiment. The fine texture had image blobs with a size of 75 µm, the coarse texture had blobs of 300 µm. In adult zebrafish, this equates to approximately 3.5° and 14.3° of retinal surface for fine and coarse texture, respectively [49]. The pixel intensity of the textures ranged from 30-90 with a mean intensity of 60 on the 8-bit DLP projector scale (0-255).

In order to simulate saccades, the texture was rapidly displaced for 100 ms at 100 to 300°/s at a straight trajectory in random direction. At various time points around the texture displacement (-150, 17, 50, 100, 250, 500, and 2000 ms after texture displacement onset) a probe flash was shown. The probe flash was a full-screen positive or negative luminance step (“bright” or “dark” flash), that lasted for one frame (16.7 ms). Local contrast within the texture remained constant between the probe flash and the background texture. In total a single trial consisted of 15 conditions (7 delays with dark flash, 7 delays with bright flash, 1 control displacement without flash) shown in random order; trials were repeated at least four times for each texture type. Additionally, the following stimuli were shown to characterize RGCs types and receptive field properties: 1. Full field contrast steps. To test ON/OFF properties of RCGs, uniform background luminance steps from 30 to 60 and from 60 to 90 pixel intensity steps in positive (ON) and negative (OFF) direction were shown. 2. Binary checkerboard flicker. For spatial receptive field mapping a binary checkerboard pattern, with checkers of 55 x 55 µm size, randomly switching between 10 or 50 pixel intensity independent of other checkers was shown. The stimulus was shown for 10 minutes. Linear filters, representing spatial and temporal receptive fields were calculated through reverse correlation.

#### Data analysis (MEA)

MEA recordings were high-pass filtered at a 500 Hz cutoff frequency using a tenth-order Butterworth filter. Spikes were sorted semi-manually using Matlab-based custom software, and spike waveforms and times were extracted according to Idrees and colleagues [6]. Single units were identified semi-manually by waveform. Waveforms within one unit were first clustered through principal component analysis and clusters were sorted based on waveform variability. Clusters with unreasonable inter-spike times or large waveform variability were excluded. RGC types were characterized based on their responses to full-field contrast steps. For each cell an ON/OFF index was calculated by dividing differences between ON and OFF step peak response by their sum. The index ranged from -1 to +1, where -1 indicates a full OFF cell and +1 indicates a full ON cell. Cells with an ON/OFF index > 0 where only analyzed for bright probe flashes and cells with an ON/OFF index < 0 were only analyzed for dark probe flashes. Further, we characterized the spatial receptive field of each cell, by examining the linear filters for each region (checker) defined by the binary checkerboard stimulus. A 500 ms window prior to each spike was summed and the modulation strength of each linear filter, measured as the standard deviation along the 500 ms kernel, determined how strongly that checker drives the RGC. The resulting 2D map of standard deviation values was fitted with a 2D Gaussian, and receptive field size was determined as an ellipse with 1.5*σ*. Receptive field sizes were converted from *x* μm to *α* degrees approximating the focal length (*f*) of the adult zebrafish eye as 1.2mm [49], such that 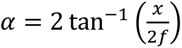. Response latency was determined as the time from stimulus onset to 90% peak response. All spike times were converted into firing rates (spikes/s) by convolving spike times with a Gaussian of *σ* = 10 ms standard deviation and amplitude 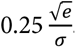.

To determine retinal saccadic suppression for each recorded RGC, first baseline responses were defined as peak firing rates in response to probe flashes at 2000 ms delay from texture displacement onset, averaged across trials. Then, average peak firing rates of the same RGC to all other delay times were normalized to this baseline response. Any texture displacement related responses were identified in control trials, without probe flashes and averaged across trials for coarse and fine textures independently. Flash responses were isolated from texture displacement responses by subtracting average texture displacement responses from the total responses, leaving only the flash-related response.

### *In-vivo* calcium imaging

#### Visual stimulation and recording setup

All visual stimuli used in this study were displayed on a custom-built spherical stimulation arena, with simultaneous eye position recordings. In the stimulation arena, fish were placed inside a custom printed Ø 80 mm resin bowl (Accura ClearVue, SLA resin, 3D Systems GmbH, Meorfelden-Walldorf, Germany), evenly coated in white acrylic spray paint and sealed with clear varnish. The resin fish bowl served as a projection surface for a DMD video projector (LightCrafter E4500 MKII, EKB Technologies Ltd., Israel), powered by a multispectral high power LED light source (CHROLIS, 365/420/470/625 nm, Thorlabs GmbH, Bergkirchen, Germany). In order to cover the entire projection surface of the bowl, the image of the video projector was split into four paths by a pyramidal mirror and refocused onto the fish bowl from all four sides (Fig. 4A) to cover -90° to +45° in elevation and 360° in azimuth. All stimuli were presented at 60 Hz refresh rate at 625 nm (red) to avoid light interference with calcium imaging. Irradiance and corresponding Michelson contrast inside the fishbowl were measured using an optical power meter (Newport 1918-R optical power meter with 918D-SL-OD3R detector, Newport Corporation, Irvine, CA, USA) and a Ø 5 mm liquid core light guide (Thorlabs GmbH, Bergkirchen, Germany) positioned on the inside wall of one half bowl at various elevations. At 625 nm stimulation wavelength the following measurements for high irradiance, low irradiance and Michelson contrast were made at 22.5°, -22.5°, and -77.5° elevation: 22.5°: 8.56 mW/m^2^ high, 0.51 mW/m^2^ low, 0.89 contrast; -22.5°: 17.83 mW/m^2^ high, 0.61 mW/m^2^ low, 0.93 contrast; -77.5°: 5.79 mW/m^2^ high, 0.61 mW/m^2^ low, 0.81 contrast.

A custom-built 850 nm IR-LED ring provided illumination for eye position tracking through the excitation path. A small window in the spherical stimulation arena below the fish (∼5 mm diameter) allowed for recording of eye potion with an infrared-filtered camera at 100 Hz (Basler ace 2 R, a2A1920-160umBAS, Basler AG, Ahrensburg, Germany). *VxPy* (Version 0.1.4) was used to generate and display visual stimuli, as well as record eye position traces and trigger visual stimuli in a closed-loop manner.

#### Visual stimulation protocol (Calcium Imaging)

Visual stimuli were designed in analogy to retinal stimulation, but adjusted to the geometry of our stimulation arena. Instead of using a rectangular array of black and white pixels and convolving these with Gaussians, we designed a stimulus in spherical coordinates by first placing equidistant nodes on the surface of a unit sphere. Half of the nodes were randomly selected as “bright” nodes, and identical von-Mises-Fisher distributions with concentration, κ= 5.6 were placed centered on each of these. Nothing was placed on the remaining, “dark” nodes. After summing up all distributions, intensities were rescaled to impose a desired minimum and maximum across the sphere surface to ensure roughly uniform local luminance and contrast across the visual field. The number of nodes is the parameter that determines pairwise distances between them. Its values were chosen to produce stimuli with spatial frequencies (Fig. S9) that reliably elicit behavioral responses. For the coarse texture 2,000 nodes (1,000 bright) were used, for the fine texture 4,000 nodes (2,000 bright) were used. In “texture displacement” experiments, saccades were simulated through a rapid rotation of the texture on the vertical axis by 30° at 300°/s either to the left or right. In “real saccade” experiments, saccades were detected with a velocity threshold of at least 200°/s. Saccade threshold was individually adjusted for each fish, to ensure best saccade detection, without noise interference.

At various delay times around texture displacement onset or saccade detection a probe stimulus was presented. Delay times were usually -500, 20, 100, 250, 500, 1000, 2000, and 4000 ms, unless stated otherwise. In “real saccade” experiments, no pre-saccadic stimulus could be shown. Probe stimuli were shown twice without texture displacement or 8 s after real saccade onset and used as baseline stimuli. Additionally, texture displacement/saccade occurred without probe stimulus. Therefore, one trial consisted of 9-11 conditions (6-8 delays, 2 baseline probes, 1 control displacement/saccade), shown in random order; trials were repeated at least 4 times.

In flash experiments the secondary visual stimulus consisted of a whole-field luminance decrease with a cosine shape, lasting 500 ms at 0.5 Hz and an amplitude of 50% luminance change. For high frequency recordings, a luminance step, lasting 20 ms with an amplitude of 50% luminance decrease was used.

In moving dot experiments, a dark dot, moving across the static texture was used as a secondary visual stimulus. The moving dot had a 20° diameter and was located in the upper left visual field at 10° elevation. The dot moved at 60°/s in azimuth from -30° to -60° and back to 0° over a time span of 1.5 s. To avoid movement overlap between the texture displacement (lasting 100 ms) and the moving dot stimulus, only delay times ≥ 100 ms were used in the moving dot experiment.

In looming disc experiments a dark disc expanded in diameter (θ) from the south pole upward with a starting diameter of *θ_0_* = 4° and a *l/v* ratio of 120 ms according to the following expansion formula: 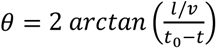, where 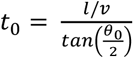 is the time of collision, until a final size of 180° was reached [29,50]. Here, the stimulus halted and remained until 5 sec had passed since looming onset. Here, also only delay times ≥100 ms were used.

#### Two-photon calcium imaging

For in-vivo calcium imaging, fish were embedded in 1.6% low-melting agarose and placed onto a custom-built stage, which could be attached to a holder on the inside of the spherical stimulation arena. For the stage, the tip of a plastic Pasteur pipette was cut and shaped into an arrowhead shape. Then, the pipette tip was placed into a custom 3D-printed mold and 2.5% low-melting agarose was poured into the mold resulting in a transparent stage that allowed the fish to be placed central inside the spherical stimulation arena. For “real saccade” experiments, agarose was removed around the eyes of the fish, to allow for free eye movements. Once the fish was placed in the center of the spherical stimulation arena dorsal side up, it was filled with clear E3 medium. The position of the fish could be adjusted, to correct for potential roll.

Calcium activity of tectal neurons was recorded with a movable objective two-photon setup (Sutter Instruments, Novato, California, USA), previously described [51]. The calcium indicator was excited at 920 nm and emitted light was collected with a 20x/1.0 Zeiss objective. In most recordings image acquisition rate was 2.18 fps, with 512 x 512 pixels imaged. Only in high-speed recordings a smaller window (40 x 40 pixels) was imaged at 50 fps acquisition rate. Magnification ranged from 1-2x, where 1x results in a resolution of 0.88 µm x 0.88 µm per pixel.

#### Data analysis (Calcium imaging)

Calcium recordings with 2.18 Hz acquisition rate were registered and segmented with suite2p [52] into individual ROIs, each representing one tectal neuron. A time constant of 1.61 s was used and for recordings at 1.6x magnification a cell diameter of 10 pixels (5.5 μm) was used. For different magnifications, the cell size was adjusted accordingly. The suite2p built-in selection algorithm *is-cell* was ignored. High-speed recordings with 50 fps acquisition rate were segmented manually in Fiji (ImageJ) into elliptical ROIs, based on standard deviation projections. Center positions of segmented ROIs were registered to the Max Planck Zebrafish Brain (mapzebrain) atlas [53] using ANTsPy [54].

Further data analysis was performed using custom-written Matlab (The MathWorks Inc., Natick, USA) and Python (PyCharm, JetBrains, Prague, Czech Republic) scripts. Fluorescence traces for each ROI were normalized into ΔF/F_0_, with a sliding window median of 120 seconds for F_0_. Through a regression-based analysis, ROIs that responded with high reproducibility across trials to the secondary visual stimulus used in the experiment (flash or moving dot), when shown in temporal isolation (“baseline”, i.e. without saccade or texture displacement), were identified. For that, calcium traces 5s prior and 10s after onset of “baseline” stimuli were averaged across trials and correlated to a calcium impulse response function, according to the calcium indicator dynamics. Further, “baseline” calcium traces were auto-correlated across trials to test for reproducibility. All ROIs with a positive significant regressor correlation and an average auto-correlation > 0.2 were considered for further analysis (4,924/63,984 ROIs across all flash experiments, 7.70%, 956/12,256 ROIs for moving dot experiment, 7.80%, 1,278/23,897 ROIs for looming disc, 5.35%).

Saccade- and texture displacement-associated activity in ROIs was identified, by averaging calcium traces during the saccade/texture displacement from control conditions (without secondary visual stimulus) and long delay conditions (≥2000 ms) up to the point of secondary visual stimulus presentation. Average saccade-/texture displacement-associated activity was then subtracted from the calcium trace for the time points during and shortly after the saccade/texture displacement.

Peak ΔF/F responses to the secondary visual stimulus at each delay time were quantified and averaged across trials for each ROI. For “real saccade” experiments, trials in which fish performed undesired saccades between the initial saccade and presentation of the secondary visual stimulus were discarded. Trials, in which the fish struggled, thus rapidly moved its head, rather than performing an actual saccade were also discarded. Average peak ΔF/F responses were normalized to average baseline responses without saccade/texture displacement. Individual modulation curves for each ROI were generated as in Figs. 4C and S4B.

To identify suppressed ROIs, amongst all ROIs sensitive to the visual stimulus, modulation curves of individual ROIs were first correlated against 6-8 model suppression curves (Fig. S4C), depending on the number of delays used in the experiment. Model suppression curves had different delay times of peak suppression and each a unimodal modulation. The average modulation across all delays had to be <1, thus suppressed. Out of the model suppression curves, the curve with the best correlation coefficient was then confirmed through a permutation test with a Bonferroni-corrected (n = 6-8) significance cut-off of p < 0.05. ROIs that passed that permutation test were considered truly suppressed (Fig. S4).

### Behavioral assay

#### Recording setup (behavioral assay)

Behavioral experiments were performed with groups of 10 freely swimming larval zebrafish placed in E3 medium in a custom 3D-printed dish. The dish was designed with inward-tapered walls aligned to the camera’s field of view to prevent visibility (of the now inclined) wall in the top-down recordings and to reduce mirror-image artefacts at the dish boundary, thereby improving downstream posture tracking. The dish was positioned centrally inside a custom-built visual stimulation arena composed of four LCD screens arranged in a square facing inward (Fig. 1A). Visual stimuli were generated and presented at 60 Hz refresh rate using *VxPy* (Version 0.1.4). Below the arena a custom-built 850 nm IR light source illuminated the animals to allow for posture tracking with an infrared-filtered camera at 84 Hz (Basler ace 2 R, a2A1920-160umBAS, Basler AG, Ahrensburg, Germany) from above.

#### Visual stimulation protocol (behavioral essay)

Free-swimming animals were presented with a coarse texture as described in the calcium imaging experiments. To simulate saccades, the texture was displaced horizontally for 100 ms at 30°/s. At delay times of -500, 20, 100, 250, 500, 1000, 2000, and 4000 ms from texture displacement onset, a global luminance step (dark-flash) with a 30% decrease in luminance lasting 500 ms was presented. Texture displacements were shown without dark-flash and dark-flash was also shown without texture displacement (baseline condition). Trials were repeated four times with an intertrial interval of 60 sec.

#### Data analysis (behavioral assay)

All videos were preprocessed by applying a static spatial mask around the dish boundary in Fiji (ImageJ) to exclude wall regions from analysis. Individual fish were tracked using TRex [55] in combination with a custom-trained YOLO11x pose-estimation model (Ultralytics, Frederick, MD, USA). The pose model was trained on a combined dataset consisting of publicly available annotated larval zebrafish images [56] and additional expert annotations. For pose estimation, 15 keypoints were defined per fish: four eye landmarks (top and bottom of each eye), one swim bladder landmark, and ten equally spaced landmarks along the midline of the tail (Fig. 1B).

O-bend events were quantified from the predicted midline geometry by computing a cumulative bending metric along the swim-bladder-to-tail axis from consecutive triplets of keypoints; events were classified as O-bends when the absolute cumulative bend exceeded 130°. For O-bend probability analyses, O-bends occurring within 1 s after probe flash onset were counted and converted into per-fish response probabilities for each flash delay condition. For O-bend latency analyses, the onset time of the first detected O-bend within 2 s after flash onset was extracted and converted into a latency relative to flash onset. Latencies were averaged per fish and delay condition. Fish were excluded from group statistics if they showed no movement, if their mean O-bend probability across stimulus conditions was < 0.1, or if they showed no O-bend responses in baseline trials (probe flash presented without texture displacement).

### Statistics

Graphical presentation and statistical analysis was done in Matlab. For retinal electrophysiology recordings, all recorded RGCs were considered in group data. For *In-vivo* calcium imaging, only truly suppressed ROIs (as described above) were considered for group data. Group data were tested for normality using a Lilliefors test. Since almost all data did not pass this test we chose non-parametric tests for further statistical analysis. Wilcoxon signed-rank tests were used for paired observations; Wilcoxon rank-sum tests were used for independent sample testing. For regression-based analysis and to identify truly suppressed ROIs in *in-vivo* calcium recordings, Pearson linear correlation was used. For correlation between saccade size and normalized flash response (Fig. 6G, Spearman rank correlation was used. A χ^2^-test for proportions was used to compare proportions of suppressed ROIs across experiments (Fig. 8B). For all tests p-value cutoffs were Bonferroni-corrected for multiple testing, when necessary and p < 0.05 was considered as significance cut-off.

## Data availability

The scripts and the required pre-processed dataset for the figures and supplementary figures included in the current study are available in a public repository (https://doi.gin.g-node.org/…) upon publication. The retinal electrophysiology data and raw calcium imaging data supporting the current study have not been deposited in a public repository because of their large size but are available from the corresponding author on request.

## Acknowledgements

We want to thank all members of the Arrenberg group for fruitful discussions regarding this work, Saad Idrees for his incredible support and expertise regarding retinal electrophysiology and we want to thank Lidia García-Pradas for her excellent fish care.

## Supporting information

**Fig. S1. Receptive field properties of zebrafish RGCs**. **(A)** Receptive field sizes and response latencies were characterized using a binary checkerboard flicker with a checker size of 55 x 55 μm. Orange ellipse indicates receptive filed size and orange line indicates response latency of an example RGCs, both determined as described in Methods. Group data (right panels) shows receptive filed sizes and response latencies of all RGCs recorded, error bars indicating mean ± SD. *N = 87 RGCs*. **(B)** Whole-field contrast steps, each lasting 2 seconds were used to determine preferred polarity of RGCs. A majority of RGCs preferred light-to-dark (OFF) contrast steps over dark-to-light (ON) contrast steps. **(C)** Spike count for individual trials and mean firing rate across trials during whole-field contrast steps for one example OFF RGC (upper panels) and one example ON RGC (lower panels).

**Fig. S2. Texture displacement and saccade induced modulation of visual responses in OT. (A)** Texture displacement-induced modulation of flash responses at various flash delays of all visually responsive ROIs for displacement of coarse texture (left panel), fine texture (middle panel), and for shuffled data (right panel). Visual responses are diversely modulated up and down. Real data shows, that more neurons are suppressed, than enhanced and enhancement is weaker, compared to what is expected by chance in shuffled data. **(B)** Saccade-induced modulation of flash responses at various flash delays of all visually responsive ROIs for saccades across textured background (left panel), saccades across uniform background (middle panel), and for shuffled data (right panel). Visual responses are more strongly modulated than by texture displacement (compare panel a.), with more suppressed than enhanced neurons in real data, compared to shuffled data. In both panels A and B, cells are sorted along the y-axis according to the following parameters: 1) The overall percent modulation across all delay times is calculated for each cell and cells are sorted in ascending order, 2) all cells are split up in 5 bins (each containing 20% of the cells), and within the bin, neurons are sorted according to the delay time with the strongest modulation (suppression or enhancement), 3) for each peak modulation delay time (within a 20% bin), the corresponding cells are sorted according to their overall suppression or enhancement across all delay times.

**Fig. S3. Anatomical distribution of saccade-modulated ROIs.** Sagittal and coronal view of center positions of all flash-responsive ROIs. Flash-responsive ROIs can be found all over the tectum, as is expected from the distribution of all registered ROIs in the tectum (p = 0.08 for enhanced and suppressed ROIs vs all ROIs on AP-axis, p = 0.62 for enhanced and suppressed ROIs vs all ROIs on ML-axis, p = 0.06 for suppressed and enhanced ROIs vs all ROIs on DV-axis, two-tailed Wilcoxon rank-sum test). Suppressed ROIs can be found more posteriorly, compared enhanced ROIs (p < 0.001 for suppressed vs enhanced ROIs on AP axis, p = 0.44 for suppressed vs enhanced on ML-axis, p = 0.99 for suppressed vs enhanced ROIs on DV-axis, two-tailed Wilcoxon rank-sum test).

**Fig. S4. Identifying suppressed ROIs. (A)** Measured peak ΔF/F responses to probe flash of sample ROI for individual trials and averaged across trials. **(B)** Mean responses of same ROI as in a. normalized to baseline responses (visual response in absence of texture displacement/saccade). **(C)** Model suppression curves with various temporal suppression dynamics. For each ROI, the best-matching model suppression curve was determined using Pearson correlation. For each ROI, the existence of true suppression was investigated using a permutation test. **(D)** Results of permutation test of same ROI as in (A) and (B). Delay identities were permuted 10,000 times and correlations to the best-matching model curve were calculated for each permuted dataset, revealing a significance cutoff of R ≥ 0.58879. The sample ROI passed the permutation test with R = 0.80826 and can be considered as significantly suppressed.

**Fig. S5. Saccadic suppression does not last indefinitely and is not due to stimulus artefacts or simple stimulus-stimulus interaction.** (**A)** Temporal suppression profile for delays > 4 sec. Flash responses return to baseline levels by 8 sec after texture displacement onset and remain unchanged for longer delays (# p < 0.0001, two-tailed Wilcoxon signed-rank test for suppression, Bonferroni corrected for n = 11 conditions, *n = 490 ROIs from 3 animals*). (**B**) The reported slow temporal suppression dynamics of flash responses are not due to slow calcium imaging or long-lasting visual stimulus. In a small subset of recordings, we tested a short 20 ms flash stimulus, comparable to the brief probe flash used in the high temporal resolution MEA recordings in the retina. To capture small and rapidly changing neural responses to this brief visual stimulus, we chose to image a much smaller brain region, but at 50 fps temporal resolution. We found that even with this brief visual stimulus, and with recordings at high temporal resolution, suppression lasting >1000 ms remained. Data shown are mean ± SEM of percent suppression for each delay time for all significantly suppressed ROIs (# p < 0.001, two-tailed Wilcoxon signed-rank test for suppression, * p < 0.05, two-tailed Wilcoxon signed-rank test for texture, Bonferroni corrected for n = 24 conditions, *n = 30 ROIs from 6 animals*). (**C**) While some part of saccadic suppression can be explained by simple stimulus-stimulus interaction, pairing two identical probe flashes (red) or one short 20ms flash, representing a short visual event, such as a saccade, with a probe flash, leads only to very small (<5%) suppression for up to 1000 ms from first flash onset (# p < 0.01, two-tailed Wilcoxon signed-rank test for suppression, Bonferroni corrected for n = 10 conditions, *n = 610 ROIs from 3 animals*).

**Fig. S6. Saccade size and flash response modulation.** For early delays <500ms, saccade size negatively correlates with normalized flash response, where larger saccades across coarse or fine texture led to more strongly suppressed flash responses (p = 0.0036, Spearman rank correlation, n = 284 saccades for coarse, <500ms, p < 0.001, Spearman rank correlation, n = 259 saccades for fine, <500ms). For late delays ≥500 ms, there was no correlation between saccade size and suppression for both textures (p = 0.84, Spearman rank correlation, n = 400 saccades for coarse, ≥500ms, p = 0.26, Spearman rank correlation, n = 387 saccades for fine, ≥500ms).

**Fig. S7. Anatomical distribution of moving dot and looming disc associated ROIs. (A)** Dorsal, sagittal and coronal view of center positions of all moving dot-responsive ROIs. Moving dot-responsive ROIs (pink line) were located more medially and more anteriorly, compared to all ROIs (grey line) that could be segmented within the recording window (p < 0.0001 for ML-axis, p < 0.0001 for AP-axis, two-tailed Wilcoxon rank-sum test, Bonferroni corrected, n = 327 dot-responsive ROIs, 4,746 total registered ROIs). (**B**) Dorsal, sagittal and coronal view of center positions of all looming disc-responsive ROIs. Looming disc-responsive ROIs (green) were located more ventrally, compared to all ROIs (grey) that could be segmented within the recording window (p < 0.0001 for DV-axis, two-tailed Wilcoxon rank-sum test, Bonferroni corrected, n = 1,823 loom-responsive ROIs, 23,820 total registered ROIs).

**Fig. S8. Distribution of flash, moving dot and looming responses.** Average responses to global flash on coarse-textured or uniform background of all flash-responsive ROIs (blue bars), and average responses to moving dot on coarse-textured background of all dot-responsive ROIs (pink bars) did not differ in size (p = 0.83, two-tailed Wilcoxon rank-sum test, n = 4,924 flash-responsive ROIs, n = 956 dot-responsive ROIs). Average responses to looming disc on coarse-textured background of all loom-responsive ROIs were larger than both flash and dot responses (p < 0.0001 for disc v. flash and disc vs. dot, two-tailed Wilcoxon rank-sum test, n = 1,879 loom-responsive ROIs).

**Fig. S9. Comparison of visual stimuli used in tectal and retinal recordings. (A)** Mercator plot of the fine (left) and coarse (right) textures presented in tectal recordings (top two panels) and retinal recordings (bottom two panels), adapted from Idrees et al. [6], shown here in visual field coordinates. Color bar and axis labels apply to all four patterns equally. Red, orange, blue and teal markers show pixels whose intensities were analyzed for subsequent panels. **(B-E)** Representative spatial frequency spectra for each of the stimulus patterns. **(B)** fine pattern for tectal recording, **(C)** coarse pattern for tectal recording, **(D)** fine pattern for retinal recording, **(E)** coarse pattern for retinal recording. For mathematical convenience, statistics were calculated along one-dimensional paths only. Because the random processes used to generate the patterns are not biased towards any direction in visual field space, these paths provide a reasonable estimate without loss of generality. Otherwise identical features covering identical solid angles will cover a different range of the azimuth if they appear at different elevations; this is accounted for by adjusting the sampling to the inverse of the sine of the elevation, with the densest sampling around the equator.

## References

1. Zuber BL, Stark L. Saccadic suppression: Elevation of visual threshold associated with saccadic eye movements. Exp Neurol. 1966;16: 65–79. doi:10.1016/0014-4886(66)90087-2

2. Beeler GW. Visual threshold changes resulting from spontaneous saccadic eye movements. Vision Res. 1967;7: 769–775. doi:10.1016/0042-6989(67)90039-9

3. Burr DC, Morrone MC, Ross J. Selective suppression of the magnocellular visual pathway during saccadic eye movements. Nature 1994 371:6497. 1994;371: 511–513. doi:10.1038/371511a0

4. Bremmer F, Kubischik M, Hoffmann KP, Krekelberg B. Neural dynamics of saccadic suppression. Journal of Neuroscience. 2009;29: 12374–12383. doi:10.1523/JNEUROSCI.2908-09.2009

5. Hafed ZM, Krauzlis RJ. Microsaccadic suppression of visual bursts in the primate superior colliculus. Journal of Neuroscience. 2010;30: 9542–9547. doi:10.1523/JNEUROSCI.1137-10.2010

6. Idrees S, Baumann MP, Franke F, Münch TA, Hafed ZM. Perceptual saccadic suppression starts in the retina. Nat Commun. 2020;11: 1–19. doi:10.1038/s41467-020-15890-w

7. Easter SS, Gregory Nicola JN. The Development of Eye Movements in the Zebrafish (Danio rerio). Dev Psychobiol. 1997;31: 267–276. doi:10.1002/(SICI)1098-2302(199712)31:4<267::AID-DEV4>3.0.CO;2-P

8. Schoonheim PJ, Arrenberg AB, Del Bene F, Baier H. Optogenetic localization and genetic perturbation of saccade-generating neurons in Zebrafish. Journal of Neuroscience. 2010;30: 7111–7120. doi:10.1523/JNEUROSCI.5193-09.2010

9. Bollmann JH. The Zebrafish Visual System: From Circuits to Behavior. https://doi.org/101146/annurev-vision-091718-014723. 2019;5: 269–293. doi:10.1146/ANNUREV-VISION-091718-014723

10. Baumann MP, Idrees S, Münch T, Hafed Z. Selective peri-saccadic suppression of low spatial frequencies is a visual phenomenon. J Vis. 2019;19: 253–253. doi:10.1167/19.10.253

11. Ridder WH, Tomlinson A. A comparison of saccadic and blink suppression in normal observers. Vision Res. 1997;37: 3171–3179. doi:10.1016/S0042-6989(97)00110-7

12. Chen CY, Hafed ZM. A neural locus for spatial-frequency specific saccadic suppression in visual-motor neurons of the primate superior colliculus. J Neurophysiol. 2017;117: 1657–1673. doi:10.1152/jn.00911.2016

13. Michels L, Lappe M. Contrast dependency of saccadic compression and suppression. Vision Res. 2004;44: 2327–2336. doi:10.1016/J.VISRES.2004.05.008

14. Knöll J, Binda P, Concetta Morrone M, Bremmer F. Spatiotemporal profile of peri-saccadic contrast sensitivity. J Vis. 2011;11: 15–15. doi:10.1167/11.14.15

15. Bianco IH, Kampff AR, Engert F. Prey capture behavior evoked by simple visual stimuli in larval zebrafish. Front Syst Neurosci. 2011;5: 18496. doi:10.3389/FNSYS.2011.00101/ABSTRACT

16. Sperling G. Temporal and Spatial Visual Masking. I. Masking by Impulse Flashes. JOSA, Vol 55, Issue 5, pp 541–559. 1965;55: 541–559. doi:10.1364/JOSA.55.000541

17. Matin E, Clymer AB, Matin L. Metacontrast and Saccadic Suppression. Science (1979). 1972;178: 179–182. doi:10.1126/SCIENCE.178.4057.179

18. Campbell FW, Wurtz RH. Saccadic omission: Why we do not see a grey-out during a saccadic eye movement. Vision Res. 1978;18: 1297–1303. doi:10.1016/0042-6989(78)90219-5

19. Idrees S, Baumann MP, Korympidou MM, Schubert T, Kling A, Franke K, et al. Suppression without inhibition: how retinal computation contributes to saccadic suppression. Communications Biology 2022 5:1. 2022;5: 1–23. doi:10.1038/s42003-022-03526-2

20. Diamond MR, Ross J, Morrone MC. Extraretinal control of saccadic suppression. J Neurosci. 2000;20: 3449–55.

21. Ali MA, Lischka K, Preuss SJ, Trivedi CA, Bollmann JH. A synaptic corollary discharge signal suppresses midbrain visual processing during saccade-like locomotion. Nature Communications 2023 14:1. 2023;14: 1–18. doi:10.1038/s41467-023-43255-6

22. Kennard DW, Hartmann RW, Kraft DP, Boshes B. Perceptual suppression of afterimages. Vision Res. 1970;10: 575–585. doi:10.1016/0042-6989(70)90051-9

23. Volkmann FC. Human visual suppression. Vision Res. 1986;26: 1401–1416. doi:10.1016/0042-6989(86)90164-1

24. Kayama Y, Riso RR, Bartlett JR, Doty RW. Luxotonic responses of units in macaque striate cortex. https://doi.org/101152/jn19794261495. 1979;42: 1495–1517. doi:10.1152/JN.1979.42.6.1495

25. Ibbotson MR, Price NSC, Crowder NA, Ono S, Mustari MJ. Enhanced Motion Sensitivity Follows Saccadic Suppression in the Superior Temporal Sulcus of the Macaque Cortex. Cerebral Cortex. 2007;17: 1129–1138. doi:10.1093/CERCOR/BHL022

26. Gu XJ, Hu M, Li B, Hu XT. The Role of Contrast Adaptation in Saccadic Suppression in Humans. PLoS One. 2014;9: e86542. doi:10.1371/JOURNAL.PONE.0086542

27. Dunn TW, Gebhardt C, Naumann EA, Riegler C, Ahrens MB, Engert F, et al. Neural circuits underlying visually evoked escapes in larval zebrafish. Neuron. 2016;89: 613. doi:10.1016/J.NEURON.2015.12.021

28. Burgess HA, Granato M. Modulation of locomotor activity in larval zebrafish during light adaptation. Journal of Experimental Biology. 2007;210: 2526–2539. doi:10.1242/JEB.003939

29. Fotowat H, Engert F. Neural circuits underlying habituation of visually evoked escape behaviors in larval zebrafish. Elife. 2023;12. doi:10.7554/ELIFE.82916

30. Huang KH, Ahrens MB, Dunn TW, Engert F. Spinal Projection Neurons Control Turning Behaviors in Zebrafish. Current Biology. 2013;23: 1566–1573. doi:10.1016/J.CUB.2013.06.044

31. Easter SS, Nicola GN. The development of vision in the zebrafish (Danio rerio). Dev Biol. 1996;180: 646–663. doi:10.1006/dbio.1996.0335

32. Dehmelt FA, Meier R, Hinz J, Yoshimatsu T, Simacek CA, Huang R, et al. Spherical arena reveals optokinetic response tuning to stimulus location, size, and frequency across entire visual field of larval zebrafish. Elife. 2021;10. doi:10.7554/ELIFE.63355

33. Dowell CK, Lau JYN, Bianco IH. The saccadic repertoire of larval zebrafish reveals kinematically distinct saccades that are used in specific behavioural contexts. bioRxiv. 2023; 2023.11.07.565345. doi:10.1101/2023.11.07.565345

34. Dera J, Stramski D. Maximum effects of sunlight focusing under a wind-disturbed sea surface. Oceanologia. 1986;23.

35. Wang K, Hinz J, Zhang Y, Thiele TR, Arrenberg AB. Parallel Channels for Motion Feature Extraction in the Pretectum and Tectum of Larval Zebrafish. Cell Rep. 2020;30: 442–453.e6. doi:10.1016/j.celrep.2019.12.031

36. Crapse TB, Sommer MA. Corollary discharge across the animal kingdom. Nat Rev Neurosci. 2008;9: 587. doi:10.1038/NRN2457

37. Fukutomi M, Carlson BA. A History of Corollary Discharge: Contributions of Mormyrid Weakly Electric Fish. Front Integr Neurosci. 2020;14: 561912. doi:10.3389/FNINT.2020.00042/BIBTEX

38. Cavanaugh J, Berman RA, Joiner WM, Wurtz RH. Saccadic Corollary Discharge Underlies Stable Visual Perception. Journal of Neuroscience. 2016;36: 31–42. doi:10.1523/JNEUROSCI.2054-15.2016

39. Rajkai C, Lakatos P, Chen CM, Pincze Z, Karmos G, Schroeder CE. Transient Cortical Excitation at the Onset of Visual Fixation. Cerebral Cortex. 2008;18: 200–209. doi:10.1093/CERCOR/BHM046

40. Ibbotson MR, Crowder NA, Cloherty SL, Price NSC, Mustari MJ. Saccadic Modulation of Neural Responses: Possible Roles in Saccadic Suppression, Enhancement, and Time Compression. The Journal of Neuroscience. 2008;28: 10952. doi:10.1523/JNEUROSCI.3950-08.2008

41. Cloherty SL, Mustari MJ, Rosa MGP, Ibbotson MR. Effects of saccades on visual processing in primate MSTd. Vision Res. 2010;50: 2683–2691. doi:10.1016/J.VISRES.2010.08.020

42. Hamker FH, Zirnsak M, Calow D, Lappe M. The Peri-Saccadic Perception of Objects and Space. doi:10.1371/journal.pcbi.0040031

43. Volotsky S, Vinepinsky E, Donchin O, Segev R. Long-range neural inhibition and stimulus competition in the archerfish optic tectum. J Comp Physiol A Neuroethol Sens Neural Behav Physiol. 2019;205: 537–552. doi:10.1007/S00359-019-01345-1/FIGURES/10

44. Segev R, Schneidman E, Goodhouse J, Ii MJB. Role of Eye Movements in the Retinal Code for a Size Discrimination Task. J Neurophysiol. 2007;98: 1380–1391. doi:10.1152/jn.00395.2007

45. Malevich T, Zhang T, Baumann MP, Bogadhi AR, Hafed ZM. Faster Detection of “Darks” than “Brights” by Monkey Superior Colliculus Neurons. Journal of Neuroscience. 2022;42: 9356–9371. doi:10.1523/JNEUROSCI.1489-22.2022

46. Wu W, Hafed ZM. Stronger premicrosaccadic sensitivity enhancement for dark contrasts in the primate superior colliculus. Scientific Reports 2025 15:1. 2025;15: 1–17. doi:10.1038/s41598-025-87090-9

47. Cameron DA. Mapping absorbance spectra, cone fractions, and neuronal mechanisms to photopic spectral sensitivity in the zebrafish. Vis Neurosci. 2002;19: 365–372. doi:10.1017/S0952523802192121

48. Vogalis F, Shiraki T, Kojima D, Wada Y, Nishiwaki Y, Jarvinen JLP, et al. Ectopic expression of cone-specific G-protein-coupled receptor kinase GRK7 in zebrafish rods leads to lower photosensitivity and altered responses. Journal of Physiology. 2011;589: 2321–2348. doi:10.1113/JPHYSIOL.2010.204156

49. Collery RF, Veth KN, Dubis AM, Carroll J, Link BA. Rapid, Accurate, and Non-Invasive Measurement of Zebrafish Axial Length and Other Eye Dimensions Using SD-OCT Allows Longitudinal Analysis of Myopia and Emmetropization. PLoS One. 2014;9: 110699. doi:10.1371/journal.pone.0110699

50. Temizer I, Donovan JC, Baier H, Semmelhack JL. A Visual Pathway for Looming-Evoked Escape in Larval Zebrafish. Current Biology. 2015;25: 1823–1834. doi:10.1016/J.CUB.2015.06.002

51. Zhang Y, Huang R, Nörenberg W, Arrenberg AB. A robust receptive field code for optic flow detection and decomposition during self-motion. Current Biology. 2022;32: 2505–2516.e8. doi:10.1016/J.CUB.2022.04.048

52. Pachitariu M, Stringer C, Dipoppa M, Schröder S, Rossi LF, Dalgleish H, et al. Suite2p: beyond 10,000 neurons with standard two-photon microscopy. bioRxiv. 2017; 061507. doi:10.1101/061507

53. Shainer I, Kuehn E, Laurell E, Al Kassar M, Mokayes N, Sherman S, et al. A single-cell resolution gene expression atlas of the larval zebrafish brain. 2023.

54. Avants BB, Tustison NJ, Song G, Cook PA, Klein A, Gee JC. A reproducible evaluation of ANTs similarity metric performance in brain image registration. Neuroimage. 2011;54: 2033–2044. doi:10.1016/J.NEUROIMAGE.2010.09.025

55. Walter T, Couzin ID. Trex, a fast multi-animal tracking system with markerless identi cation, and 2d estimation of posture and visual elds. Elife. 2021;10: 1–73. doi:10.7554/ELIFE.64000

56. Scholz LA, Mancienne T, Stednitz SJ, Scott EK, Lee CCY. Plug-and-Play automated behavioral tracking of zebrafish larvae with DeepLabCut and SLEAP: pre-trained networks and datasets of annotated poses. bioRxiv. 2025; 2025.06.04.657938. doi:10.1101/2025.06.04.657938

