## Supplementary Materials for "Saccadic suppression enhances saliency of ecologically relevant stimuli in the optic tectum"

Soto et al., 2026

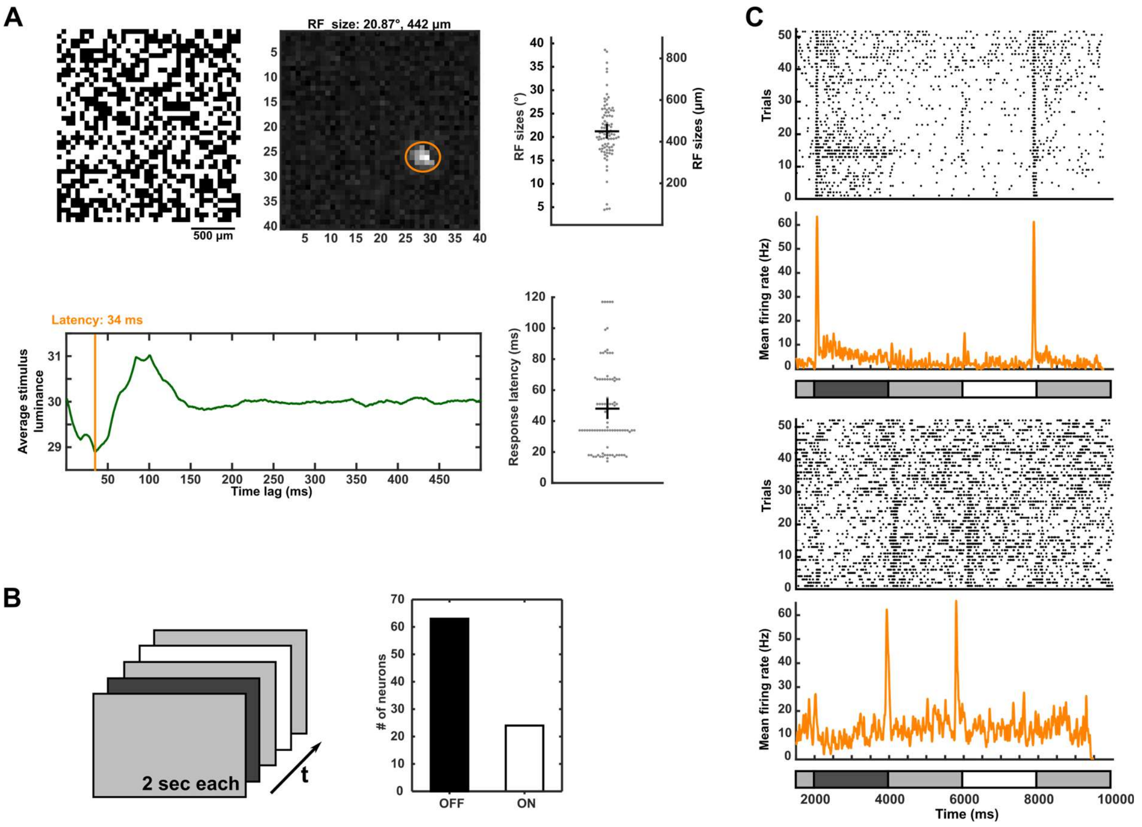


**Fig. S1. Receptive field properties of zebrafish RGCs**. **(A)** Receptive field sizes and response latencies were characterized using a binary checkerboard flicker with a checker size of 55 x 55 µm. Orange ellipse indicates receptive filed size and orange line indicates response latency of an example RGCs, both determined as described in Methods. Group data (right panels) shows receptive filed sizes and response latencies of all RGCs recorded, error bars indicating mean ± SD. *N = 87 RGCs*. **(B)** Whole-field contrast steps, each lasting 2 seconds were used to determine preferred polarity of RGCs. A majority of RGCs preferred light-to-dark (OFF) contrast steps over dark-to-light (ON) contrast steps. **(C)** Spike count for individual trials and mean firing rate across trials during whole-field contrast steps for one example OFF RGC (upper panels) and one example ON RGC (lower panels).

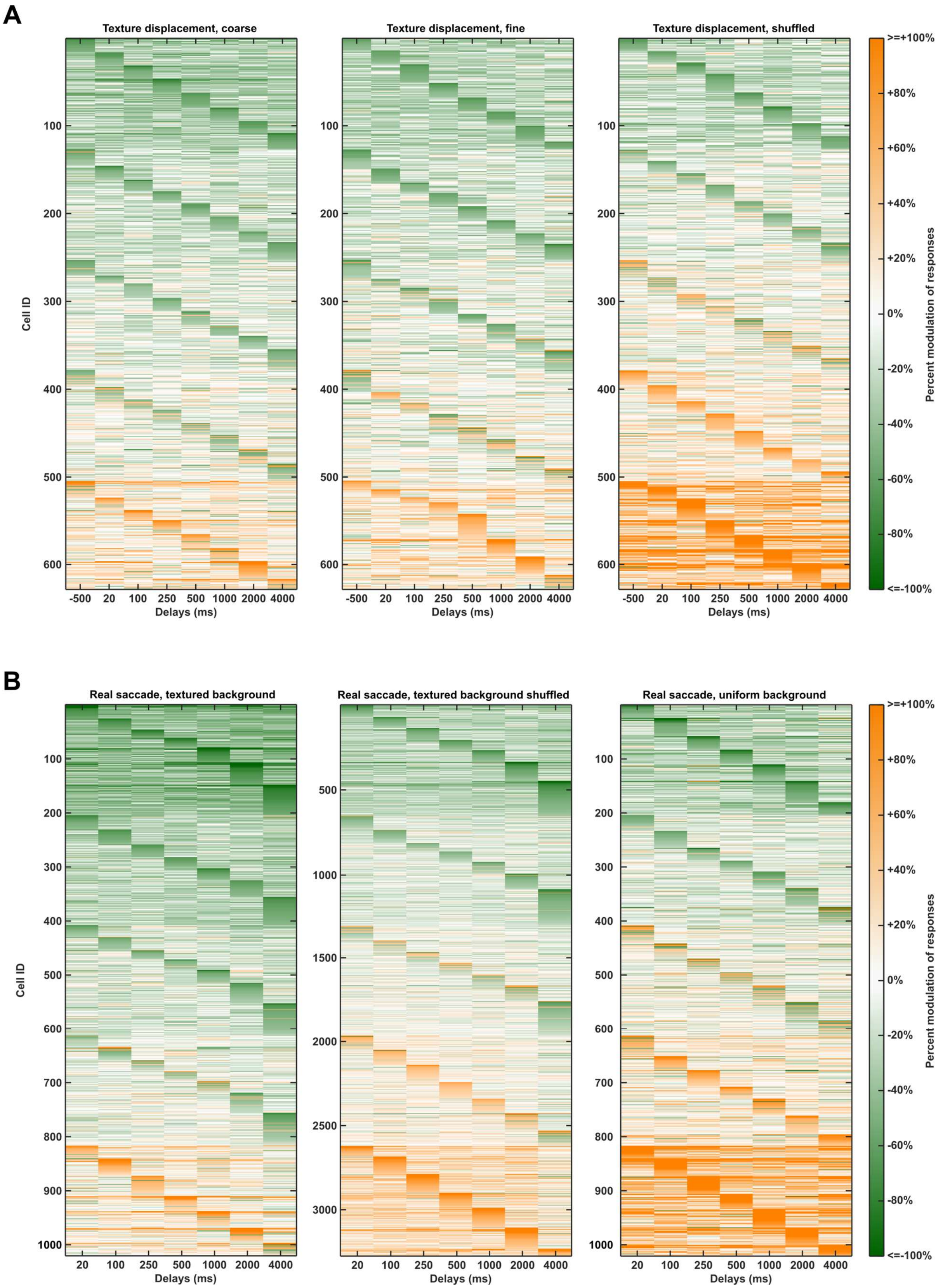

A
Texture displacement, coarse
Texture displacement, fine
Texture displacement, shuffled
Cell ID
Delays (ms)
-500 20 100 250 500 1000 2000 4000
100 200 300 400 500 600
>=+100%
+80%
+60%
+40%
+20%
0%
-20%
-40%
-60%
-80%
<=-100%
Percent modulation of responses
B
Real saccade, textured background
Real saccade, textured background shuffled
Real saccade, uniform background
Cell ID
Delays (ms)
20 100 250 500 1000 2000 4000
100 200 300 400 500 600 700 800 900 1000
500 1000 1500 2000 2500 3000
>=+100%
+80%
+60%
+40%
+20%
0%
-20%
-40%
-60%
-80%
<=-100%
Percent modulation of responses

**Fig. S2. Texture displacement and saccade induced modulation of visual responses in OT. (A)** Texture displacement-induced modulation of flash responses at various flash delays of all visually responsive ROIs for displacement of coarse texture (left panel), fine texture (middle panel), and for data with shuffled delay identities for individual trials (right panel). Visual responses are diversely modulated up and down. Real data shows, that more neurons are suppressed, than enhanced and enhancement is weaker, compared to what is expected by chance in shuffled data. **(B)** Saccade-induced modulation of flash responses at various flash delays of all visually responsive ROIs for saccades across textured background (left panel), saccades across uniform background (middle panel), and for shuffled data (right panel). Visual responses are more strongly modulated than by texture displacement (compare panel a.), with more suppressed than enhanced neurons in real data, compared to shuffled data. In both panels A and B, cells are sorted along the y-axis according to the following parameters: 1) The overall percent modulation across all delay times is calculated for each cell and cells are sorted in ascending order, 2) all cells are split up in 5 bins (each containing 20% of the cells), and within the bin, neurons are sorted according to the delay time with the strongest modulation (suppression or enhancement), 3) for each peak modulation delay time (within a 20% bin), the corresponding cells are sorted according to their overall suppression or enhancement across all delay times.

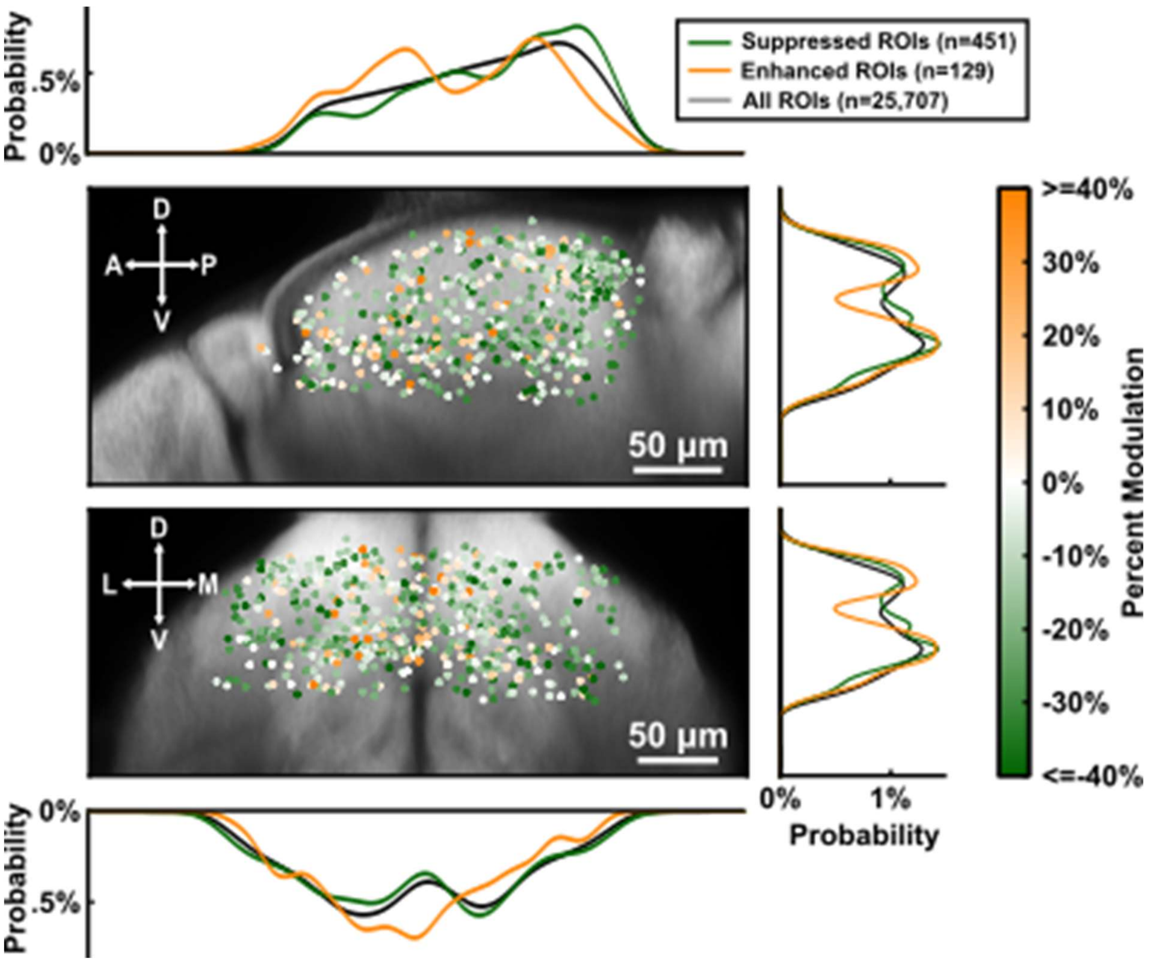


**Fig. S3. Anatomical distribution of saccade-modulated ROIs.** Sagittal and coronal view of center positions of all flash-responsive ROIs. Flash-responsive ROIs can be found all over the tectum, as is expected from the distribution of all registered ROIs in the tectum ($p = 0.08$ for enhanced and suppressed ROIs vs all ROIs on AP-axis, $p = 0.62$ for enhanced and suppressed ROIs vs all ROIs on ML-axis, $p = 0.06$ for suppressed and enhanced ROIs vs all ROIs on DV-axis, two-tailed Wilcoxon rank-sum test). Suppressed ROIs can be found more posteriorly, compared enhanced ROIs ($p < 0.001$ for suppressed vs enhanced ROIs on AP axis, $p = 0.44$ for suppressed vs enhanced on ML-axis, $p = 0.99$ for suppressed vs enhanced ROIs on DV-axis, two-tailed Wilcoxon rank-sum test).

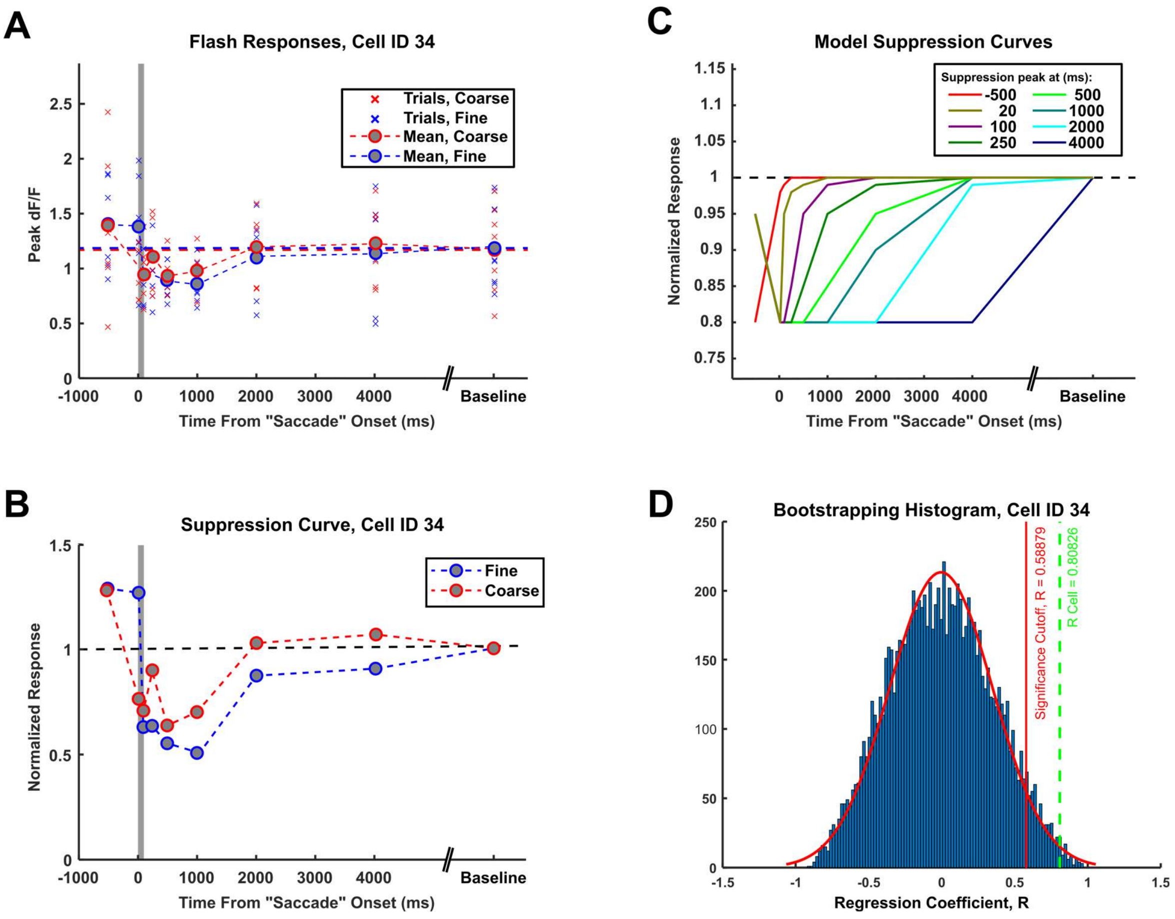


**Fig. S4. Identifying suppressed ROIs.** **(A)** Measured peak ΔF/F responses to probe flash of sample ROI for individual trials and averaged across trials. **(B)** Mean responses of same ROI as in a. normalized to baseline responses (visual response in absence of texture displacement/saccade). **(C)** Model suppression curves with various temporal suppression dynamics. For each ROI, the best-matching model suppression curve was determined using Pearson correlation. For each ROI, the existence of true suppression was investigated using a permutation test. **(D)** Results of permutation test of same ROI as in (A) and (B). Delay identities were permuted 10,000 times and correlations to the best-matching model curve were calculated for each permuted dataset, revealing a significance cutoff of $R \geq 0.58879$. The sample ROI passed the permutation test with $R = 0.80826$ and can be considered as significantly suppressed.

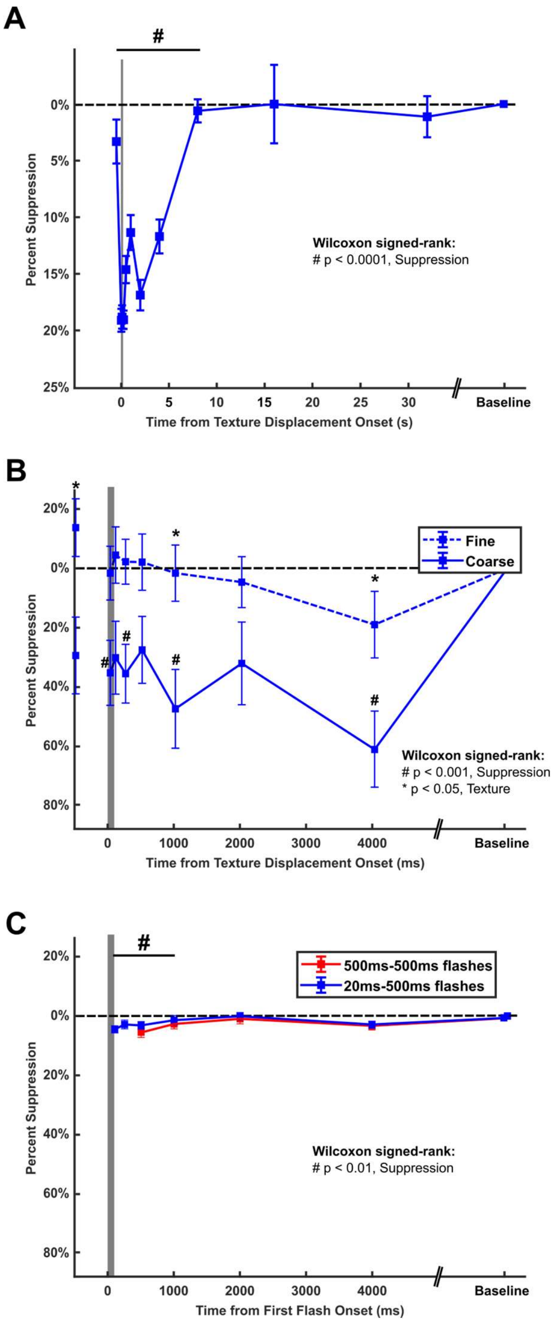


**Fig. S5. Saccadic suppression does not last indefinitely and is not due to stimulus artefacts or simple stimulus-stimulus interaction.** (**A**) Temporal suppression profile for delays > 4 sec. Flash responses return to baseline levels by 8 sec after texture displacement onset and remain unchanged for longer delays (# $p < 0.0001$, two-tailed Wilcoxon signed-rank test for suppression, Bonferroni corrected for n = 11 conditions, *n = 490 ROIs from 3 animals*). (**B**) The reported slow temporal suppression dynamics of flash responses are not due to slow calcium imaging or long-lasting visual stimulus. In a small subset of recordings, we tested a short 20 ms flash stimulus, comparable to the brief probe flash used in the high temporal resolution MEA recordings in the retina. To capture small and rapidly changing neural responses to this brief visual stimulus, we chose to image a much smaller brain region, but at 50 fps temporal resolution. We found that even with this brief visual stimulus, and with recordings at high temporal resolution, suppression lasting >1000 ms remained. Data shown are mean ± SEM of percent suppression for each delay time for all significantly suppressed ROIs (# $p < 0.001$, two-tailed Wilcoxon signed-rank test for suppression, * $p < 0.05$, two-tailed Wilcoxon signed-rank test for texture, Bonferroni corrected for n = 24 conditions, *n = 30 ROIs from 6 animals*). (**C**) While some part of saccadic suppression can be explained by simple stimulus-stimulus interaction, pairing two identical probe flashes (red) or one short 20ms flash, representing a short visual event, such as a saccade, with a probe flash, leads only to very small (<5%) suppression for up to 1000 ms from first flash onset (# $p < 0.01$, two-tailed Wilcoxon signed-rank test for suppression, Bonferroni corrected for n = 10 conditions, *n = 610 ROIs from 3 animals*).

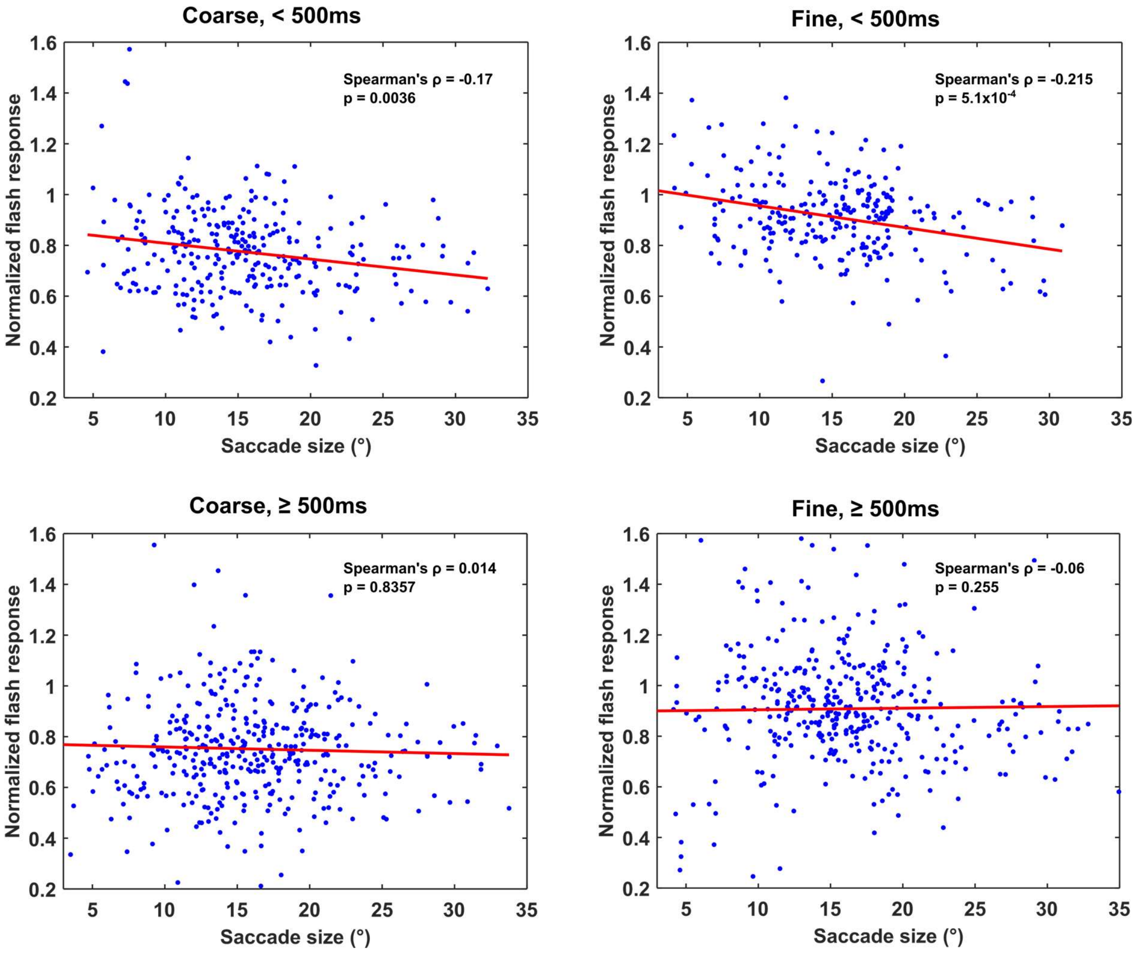


**Fig. S6. Saccade size and flash response modulation.** For early delays <500ms, saccade size negatively correlates with normalized flash response, where larger saccades across coarse or fine texture led to more strongly suppressed flash responses ($p = 0.0036$, Spearman rank correlation, $n = 284$ saccades for coarse, <500ms, $p < 0.001$, Spearman rank correlation, $n = 259$ saccades for fine, <500ms). For late delays ≥500 ms, there was no correlation between saccade size and suppression for both textures ($p = 0.84$, Spearman rank correlation, $n = 400$ saccades for coarse, ≥500ms, $p = 0.26$, Spearman rank correlation, $n = 387$ saccades for fine, ≥500ms).

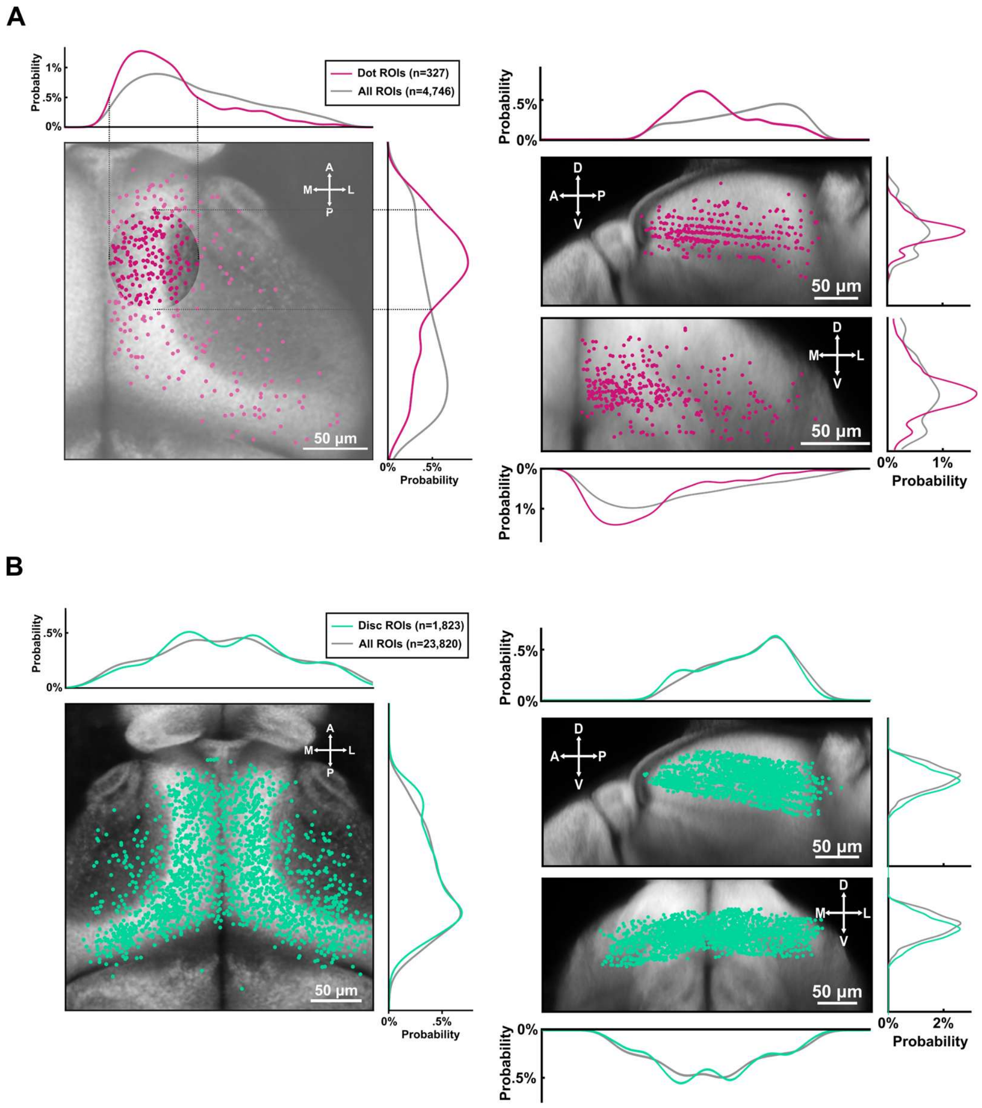


**Fig. S7. Anatomical distribution of moving dot and looming disc associated ROIs. (A)** Dorsal, sagittal and coronal view of center positions of all moving dot-responsive ROIs. Moving dot-responsive ROIs (pink line) were located more medially and more anteriorly, compared to all ROIs (grey line) that could be segmented within the recording window ($p < 0.0001$ for ML-axis, $p < 0.0001$ for AP-axis, two-tailed Wilcoxon rank-sum test, Bonferroni corrected, n = 327 dot-responsive ROIs, 4,746 total registered ROIs). (**B**) Dorsal, sagittal and coronal view of center positions of all looming disc-responsive ROIs. Looming disc-responsive ROIs (green) were located more ventrally, compared to all ROIs (grey) that could be segmented within the recording window ($p < 0.0001$ for DV-axis, two-tailed Wilcoxon rank-sum test, Bonferroni corrected, n = 1,823 loom-responsive ROIs, 23,820 total registered ROIs).

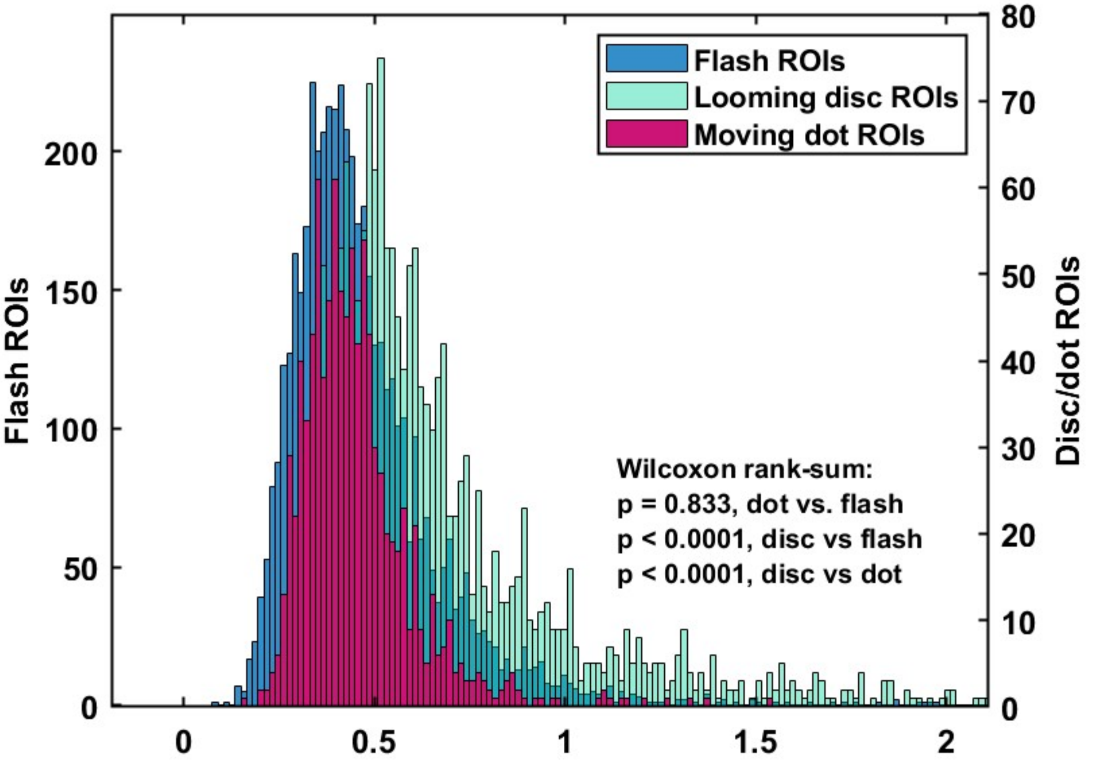


**Fig. S8. Distribution of flash, moving dot and looming responses.** Average responses to global flash on coarse-textured or uniform background of all flash-responsive ROIs (blue bars), and average responses to moving dot on coarse-textured background of all dot-responsive ROIs (pink bars) did not differ in size ($p = 0.83$, two-tailed Wilcoxon rank-sum test, $n = 4{,}924$ flash-responsive ROIs, $n = 956$ dot-responsive ROIs). Average responses to looming disc on coarse-textured background of all loom-responsive ROIs were larger than both flash and dot responses ($p < 0.0001$ for disc v. flash and disc vs. dot, two-tailed Wilcoxon rank-sum test, $n = 1{,}879$ loom-responsive ROIs).

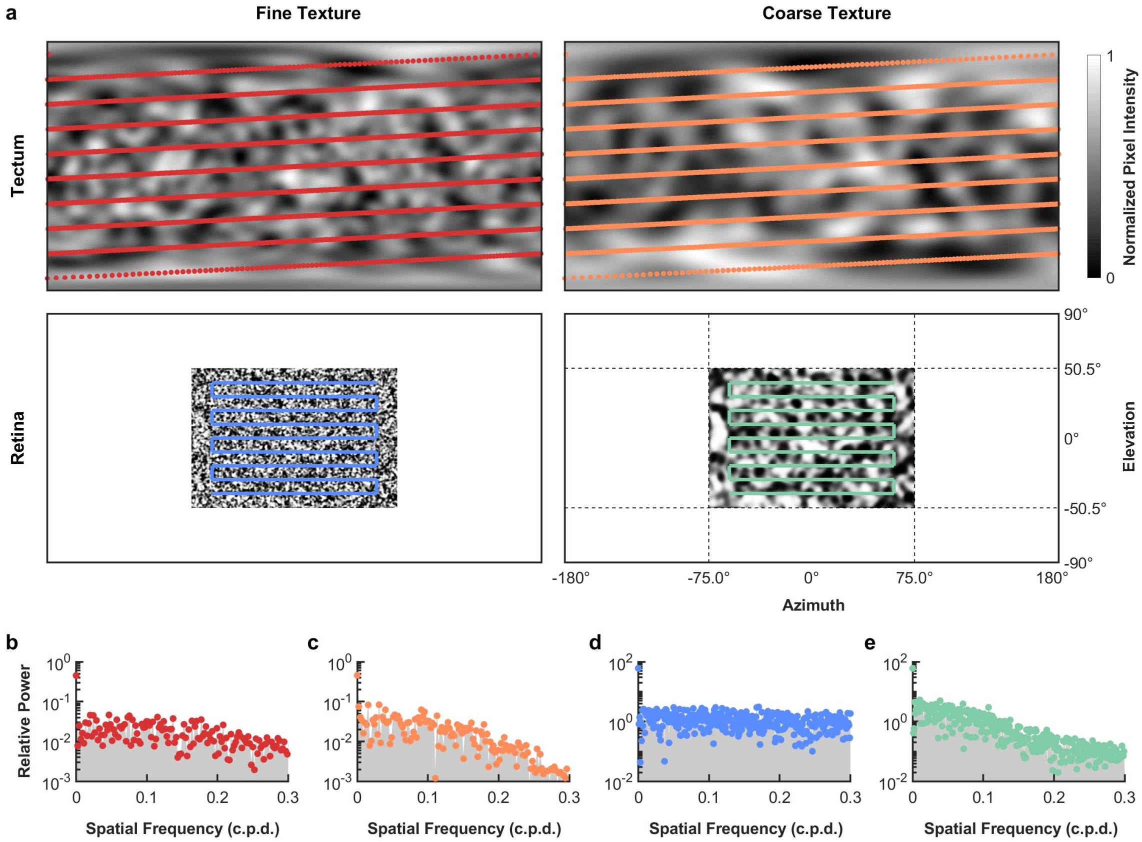


**Fig. S9. Comparison of visual stimuli used in tectal and retinal recordings. (A)** Mercator plot of the fine (left) and coarse (right) textures presented in tectal recordings (top two panels) and retinal recordings (bottom two panels), adapted from Idrees et al.[6], shown here in visual field coordinates. Color bar and axis labels apply to all four patterns equally. Red, orange, blue and teal markers show pixels whose intensities were analyzed for subsequent panels. **(B-E)** Representative spatial frequency spectra for each of the stimulus patterns. **(B)** fine pattern for tectal recording, **(C)** coarse pattern for tectal recording, **(D)** fine pattern for retinal recording, **(E)** coarse pattern for retinal recording. For mathematical convenience, statistics were calculated along one-dimensional paths only. Because the random processes used to generate the patterns are not biased towards any direction in visual field space, these paths provide a reasonable estimate without loss of generality. Otherwise identical features covering identical solid angles will cover a different range of the azimuth if they appear at different elevations; this is accounted for by adjusting the sampling to the inverse of the sine of the elevation, with the densest sampling around the equator.
